# Screening activity of brain cancer-derived factors on primary human brain pericytes

**DOI:** 10.1101/2024.09.11.612547

**Authors:** Samuel JC McCullough, Eliene Albers, Akshata Anchan, Jane Yu, Bronwen Connor, E. Scott Graham

## Abstract

Brain cancers offer poor prognoses to patients accompanied by symptoms that drastically impact the patient and their family. Brain tumours recruit local non-transformed cells to provide trophic support and immunosuppression within the tumour microenvironment, supporting tumour progression. Given the localization and supportive role of pericytes at the brain vasculature, we explored the potential for brain pericytes to contribute to the brain cancer microenvironment. To investigate this, primary brain pericytes were treated with factors commonly upregulated in brain cancers. Changes to brain pericyte cell signalling, inflammatory secretion, and phagocytosis were investigated. The TGFβ superfamily cytokines TGFβ and GDF-15 activated SMAD2/3 and inhibited C/EBP-δ, revealing a potential mechanism behind the pleiotropic action of TGFβ on brain pericytes. IL-17 induced secretion of IL-6 without activating NFκB, STAT1, SMAD2/3, or C/EBP-δ signalling pathways. IL-27 and IFNγ induced STAT1 signalling and significantly reduced pericyte phagocytosis. The remaining brain cancer-derived factors did not induce a measured response, indicating that these factors may act on other cell types or require co-stimulation with other factors to produce significant effects. Together, these findings show potential mechanisms by which brain pericytes contribute to aspects of inflammation and starts to uncover the supportive role brain pericytes may play in brain cancers.

## Introduction

Brain cancers are extremely debilitating with high mortality rates (1). A brain cancer can arise from a metastatic tumour from the body, commonly from the lung, breast, or skin (specifically melanoma), or from a transformed cell from within the brain, also known as a glioma. The most common high-grade glioma is glioblastoma (GBM), a devastating cancer with less than 50% of patients surviving two years following diagnosis (2). A necessary component of brain tumour progression is the recruitment of stromal cells such as endothelial cells, immune cells, glial cells and pericytes, which are thought to contribute close to 50% of the GBM tumour mass (3, 4). Recent studies have identified numerous tumour cell secreted factors which may modify stromal cell function, however specific pathways have yet to be identified as important regulators of stromal cell function (5, 6).

GBM, and brain-metastatic cancers like melanoma, secrete a plethora of cytokines and chemokines that influence local cells to create an ideal environment for tumour progression. These secreted factors may also be important in the recruitment of vascular cells to further promote tumour growth. Previous work in our lab has highlighted several secreted proteins from primary melanoma cell cultures which could promote tumour progression, including growth-differentiation factor-15 (GDF-15), macrophage migration inhibitory factor (MIF), platelet-derived growth factor-aa (PDGF-aa), osteopontin, and insulin-like growth factor binding protein-2 (IGFBP2) (6). Furthermore, we have identified secreted proteins from GBM cell cultures, which likely aid tumour progression, including chitinase-3-like protein 1 (CHI3L1), fractalkine, MIF, osteopontin, IGFBP2, and IGFBP3 (5). While these factors are thought to perform immune-modulatory functions, their effects on stromal cells such as brain pericytes are not known. Finally, both melanoma and GBM have been found to secrete the cytokines interleukin-17 (IL-17) and IL-27 which have been found to support (and sometimes inhibit) tumour growth (7). Given the proximity of brain pericytes to the tumour milieu and their importance in controlling the neurovascular unit, we considered the possibility that pericytes may be targets regulated by these tumour cell-secreted cytokines.

Brain pericytes can respond to numerous inflammatory cytokines, changing their function from vascular support to immunomodulatory. Pericytes secrete numerous cytokines and chemokines in response to inflammatory activation (8). Pericytes also assist in leukocyte extravasation through the expression of intercellular adhesion molecule 1 (ICAM-1) and vascular cell adhesion molecule 1 (VCAM-1) (9). Under pathological conditions, these responses can potentiate inflammatory disease through increased immune cell recruitment and activation. For example, these responses are perturbed in brain cancers whereby immune cells are recruited but become immunosuppressive, contributing to tumour immune evasion by preventing immune cell recognition of tumour antigen (10). Studies have shown that the reversal of this immunosuppressive environment in GBM results in natural killer cell-mediated killing and increased survival in mice (11). Additionally, FDA-approved immunotherapies for melanoma which target the immune checkpoint inhibitor programmed cell death protein-1 (PD-1/PD-L1) have demonstrated improved survival in patients (12). Understanding the contribution of stromal cells to mechanisms of immunosuppression in brain cancers will offer new areas of research to combat challenging diseases such as GBM, while also augmenting existing immunotherapies, such as those for metastatic melanoma.

Given the complexity of the brain cancer microenvironment, brain cancer-derived factors were added to primary human brain pericytes to elucidate the individual effects of these factors on relevant brain pericyte functions. Changes in brain pericyte cell signalling, phagocytosis, and cytokine secretion were assessed. This work highlights potential roles for brain pericytes in brain tumour progression and reveals previously unknown signalling interactions that hold significance for brain pericyte biology.

## Materials and methods

### Primary pericyte cell culture

Sciencell Human Brain Vascular Pericytes (catalogue#: 1200) were cultured using Sciencell complete pericyte medium (catalogue#: 1201) on poly-l-lysine coated flasks/plates (2µg/cm^2^) as per the manufacturer’s methods. Sciencell complete pericyte medium consists of a base medium supplemented with 2% FBS, 1% penicillin/streptomycin, and 1% pericyte growth supplement (a proprietary formulation). Poly-l-lysine was diluted to 2µg/cm^2^ in sterile milliQ water and added to the culture vessel. Coated flasks/plates were incubated at 37°C for at least 1 hour before use.

### Immunocytochemistry and inflammatory treatments

Cells were seeded at 15,000 cells/cm^2^ in 96-well plates for immunocytochemistry to detect transcription factor localisation. Cell culture samples were treated with inflammatory factors or vehicle for 1 or 4 hours (Table S1) before fixation with 4% paraformaldehyde for 15 minutes. Cells were washed with PBS and permeabilised with PBS-triton (0.1%) to reveal intracellular epitopes. Primary antibodies were diluted in immunobuffer (PBS + 1% foetal donkey serum) and were incubated with the cells at 4°C overnight (see Table S2 for details of antibody dilutions used in this study). The following day, the cells were washed 3 times with PBS before secondary antibodies diluted in immunobuffer with Hoechst 33342 were added to the cells and incubated overnight at 4°C. The following day, the stained cells were washed 3 times with PBS. Stained cells remained in PBS in a 96-well plate to be imaged.

### Cytometric bead array to study secreted factors

Cytometric bead array (CBA) experiments investigated changes in primary human brain pericyte secretion in response to inflammatory treatments. Experiments were performed as described previously (13). A list of CBA kits used in this study can be found in Table S3.

### Using fluorescent carboxylated beads to study pericyte phagocytosis

Primary human pericytes were treated with inflammatory factors for 72 hours (see Table S1), with fluorescent beads present for the final 24 hours. 50µL of Fluoresbrite YG carboxylated microspheres (1μm, catalogue#: 15702-10) diluted in pericyte medium were spiked into pericyte cell medium in 24-well plates. The beads were diluted to generate a final effector-target ratio (bead to cell ratio) of 1300:1. Cells were harvested with trypsin to cleave cell surface proteins, removing surface-bound fluorescent beads. Cells were centrifuged at 300xg for 5 minutes and resuspended in FACS buffer as a wash step (PBS, 0.5% BSA, 2nM EDTA). Brain pericytes were only washed once to conserve cell viability. The cell suspension was pelleted again (300xg, 5 mins), and the supernatant was removed before the cells were resuspended in fresh FACS buffer. Cells were stored on ice until being run on an Accuri C6 flow cytometer. Brain pericytes without bead treatment were run to show the auto-fluorescence of each pericyte population. Using the Accuri C6 Software, cellular auto-fluorescence was compared to the bead-treated pericytes to determine the percentage of phagocytic cells. The mean fluorescent intensity (MFI) of phagocytic cells was also assessed.

### Imaging methods to capture and quantify light and fluorescent images

Inflammatory transcription factors were imaged at 10x on the Molecular Devices Image Xpress Micro XLS for high-throughput screening. These images were quantified using the “Multi Wavelength Translocation” software package in MetaXpress to identify each cell as either positive or negative for a nuclear signal.

### Statistical analysis

All statistical analyses were performed using GraphPad Prism. One-way ANOVA with Tukey’s multiple comparisons determined significance between groups. EC_50_ was calculated using GraphPad Prism.

Only two repeats of each experiment were performed for factors which did not show a response, as the results between experiments corroborated to indicate the pericytes did not respond. As this work aimed to identify factors that alter brain pericyte function rather than determining specific secretory changes in response to these factors, the CBA data was generated from two repeats and statistical testing was not performed. Furthermore, given the size of the responses, there was not strong biological relevance to perform additional experiments for statistical analysis.

## Results

### Brain pericytes responded to classical inflammatory cytokines, TGFβ superfamily cytokines IL-17, and IL-27

The activity of 12 brain cancer-associated factors was screened in brain pericytes. Transcription factor translocation, phagocytosis, and cytokine/chemokine secretion were measured in response to treatment with each factor individually, as summarised in Figure 1. Transcription factor translocation was used as a broad measure of altered cellular activity. Nuclear translocation of NFκB, STAT1, and SMAD2/3 was observed 1 hour after treatment, as has been previously reported (14). Translocation of C/EBP-δ was slower than the other transcription factors, with nuclear localisation observed after 4 hours (Figure S1), in agreement with previous work (15). The activity of inflammatory factors IL-1β and TNF have been previously published but are included here as a positive control for NFκB signalling (14) (Figure 1). Of the 12 factors, 4 elicited changes to transcription factor localisation (IFNγ, TGFβ, GDF-15, IL-27). An overview of transcription factor translocation in response to each brain cancer-derived factor can be found in Table S4.

**Figure 1:**
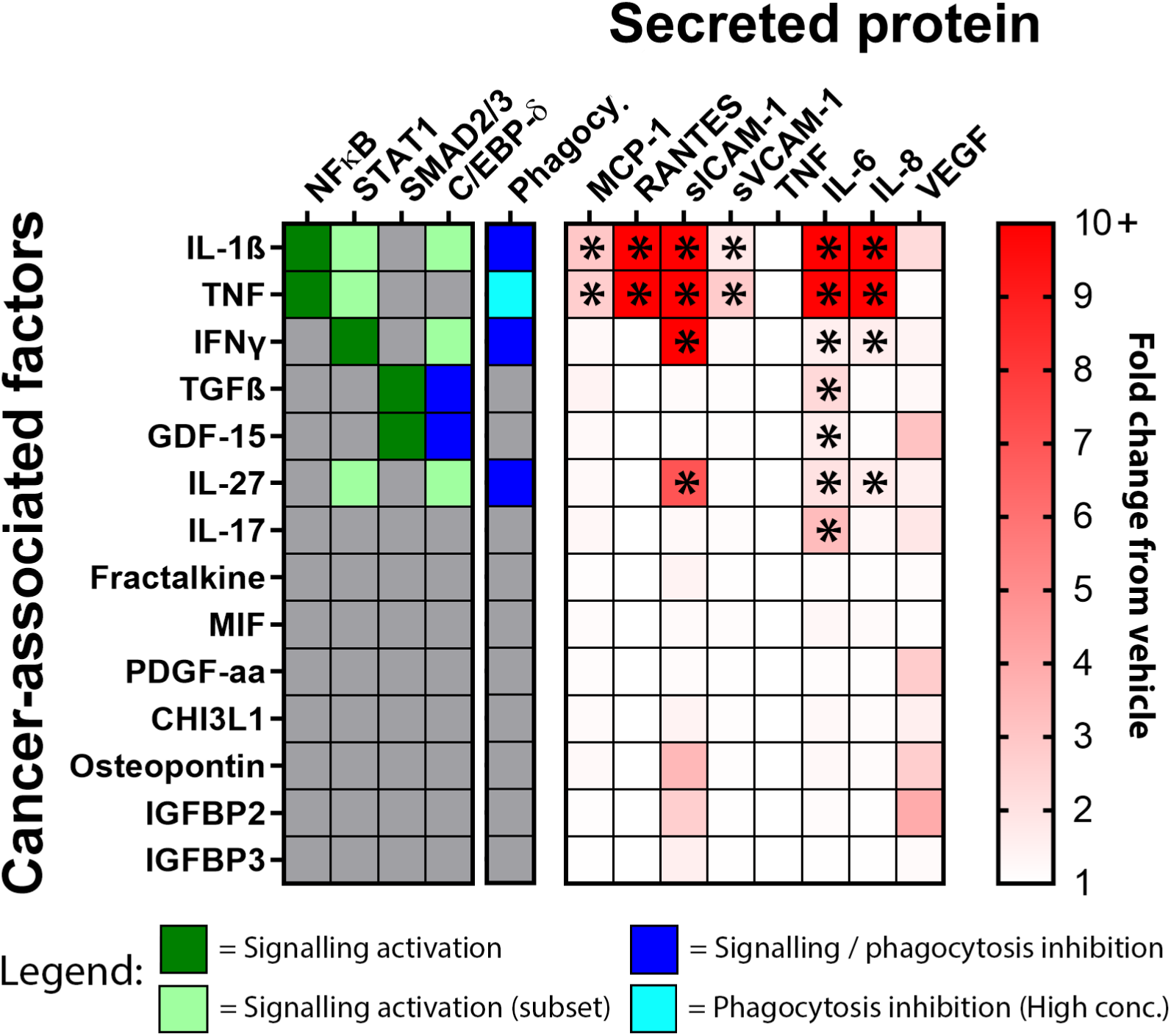
Summary of transcription factor signalling, phagocytosis, and secreted protein responses to brain cancer-associated inflammatory stimuli. The **left panel** shows a summary of brain pericyte signalling responses to each brain cancer-associated factor. Dark green indicates activation of the given signalling pathway in the majority of cultured primary pericytes. Light green indicates activation of the given signalling pathway in a minor subset of cultured primary pericytes. Blue indicates inhibition of the given signalling pathway in a subset of cultured primary pericytes. Signalling responses for NFκB, STAT1, and SMAD2/3 are measured after 1 hour, while C/EBP-δ is measured after 4 hours. The **middle panel** (named Phagocy.) shows a summary of phagocytic responses to stimuli. Dark blue indicates inhibition of phagocytosis. Light blue indicates inhibition of phagocytosis at high treatment concentrations. The **right panel** is a heat map summarising the secretory responses of primary pericytes after treatment with proteins from our panel of cancer-associated factors. The time point that demonstrated the highest fold change of secreted protein relative to vehicle is presented in the heat map for each inflammatory treatment. Observed treatment effects are denoted with * (these are not significant changes as secreted protein data is derived from 2 experimental repeats).

The 4 factors which elicited transcription factor translocation also altered the secretion of cytokines and chemokines (Figure 1). Interestingly, IL-17 treatment induced changes in secretion where no transcription factor translocation was demonstrated, suggesting that this effect is mediated by a transcription factor that was not investigated here. In general, the observed secretory changes were minor and likely did not possess biological relevance on their own. However, these changes are indicative of broad changes in brain pericyte phenotype which is worth further investigation. Changes in brain pericyte phagocytosis were also of interest as a potential mechanism of tumour immunosuppression, whereby available tumour antigen is sequestered by phagocytic pericytes to prevent immune cell recognition. Significant reductions in brain pericyte phagocytosis were demonstrated by IL-1β, TNF, IFNγ, and IL-27 (Figure 1). Treatment with fractalkine, MIF, PDGF-aa, CHI3L1, osteopontin, IGFBP2, or IGFBP3 did not induce any changes in transcription factor localisation, cytokine secretion, or phagocytosis in brain pericytes. This was unexpected given the known role these factors play in brain tumour progression. These results perhaps indicate that these factors act on different cell types or require co-stimulation to generate measurable changes in brain pericyte function (Figure 1, Figures S2-S15).

### TGFβ and GDF-15 induce SMAD2/3 signalling and inhibit C/EBP-δ activity in brain pericytes

TGFβ is an important factor for brain pericyte maturation and plays a role in regulating brain pericyte inflammation (16, 17). GDF-15 is a member of the TGFβ cytokine superfamily and is commonly upregulated in cancers due to it’s immunosuppressive functions (11). Previous studies have shown nuclear translocation of SMAD2/3 in response to TGFβ treatment in primary brain pericytes (16). The present study showed that this response is conserved in foetal primary pericytes (Figure 2). Treatment with TGFβ resulted in the concentration-dependent nuclear translocation of SMAD2/3 with a potent EC50 of 12.1pM (Figure 2A/B). On average, 68.3% of brain pericytes demonstrated SMAD2/3 translocation in response to TGFβ treatment (Figure 2A/B). Similarly, GDF-15 caused SMAD2/3 translocation in an average of 65.7% of brain pericytes, though at much higher concentrations, with an EC50 of 1515pM (Figure 2C/D). Interestingly, TGFβ inhibited nuclear translocation of C/EBP-δ with an IC50 of 15.3pM (Figure 2A/B). C/EBP-δ was inhibited in at least 17.9% of cells (Figure 2A/B). This inhibitory response was also apparent in GDF-15-treated brain pericytes, however, the concentrations used in this study were too low to produce a full concentration-response curve (Figure 2C/D). In agreement with previous studies, TGFβ and GDF-15 did not induce an NFκB or STAT1 response (14).

**Figure 2:**
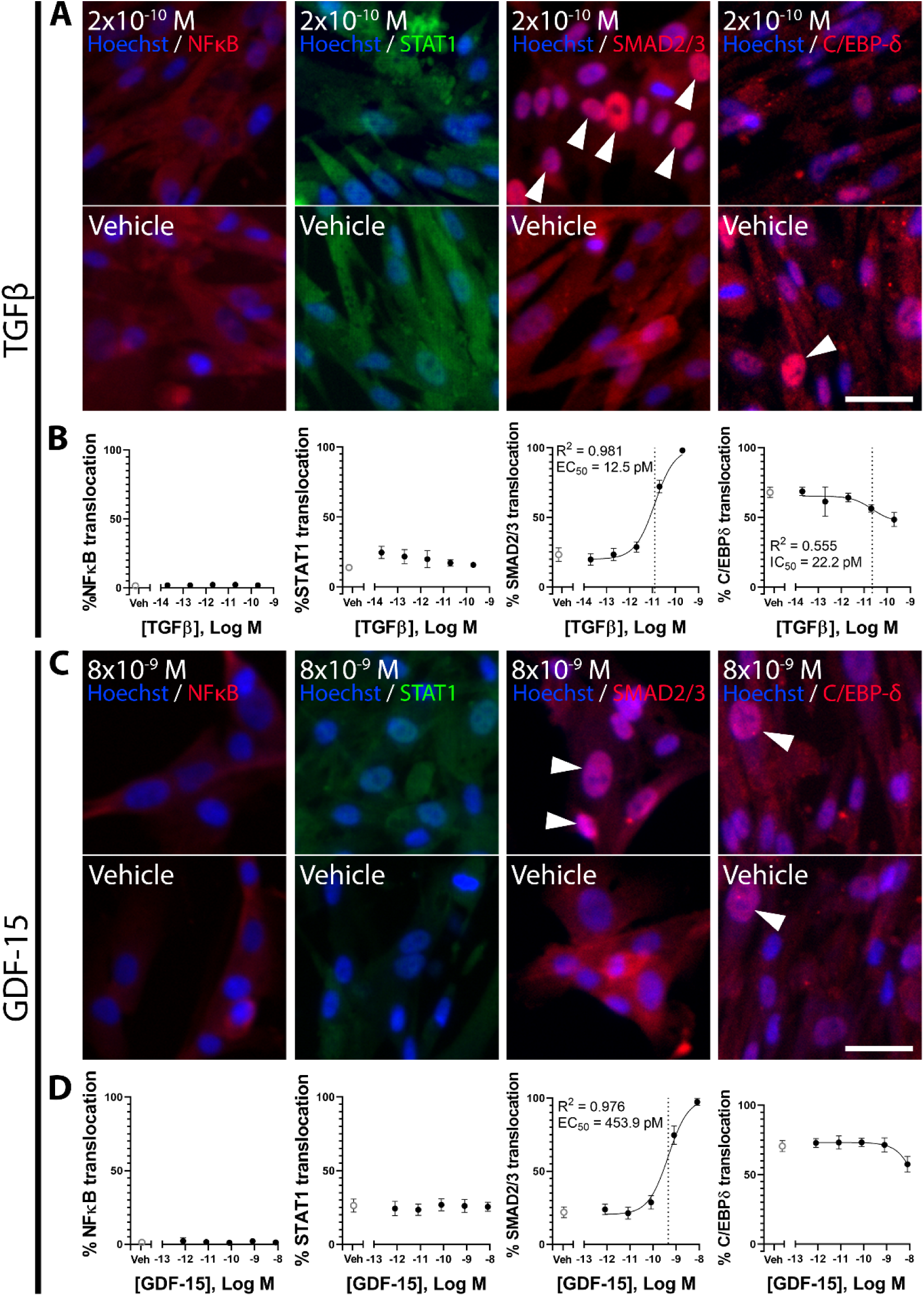
Signalling responses to treatment with TGFβ cytokine superfamily members TGFβ or GDF-15. Immunofluorescence images demonstrating NFκB, STAT1, SMAD2/3, and C/EBP-δ localisation in response to TGFβ (A) or GDF-15 (C) in primary brain pericytes. Images are quantified using MetaXpress to generate concentration-response curves (B,D). Data presented is one representative experiment of two experimental repeats for TGFβ treatment, and three experimental repeats for GDF-15 treatment (see supplemental table 4). The EC_50_ for nuclear localisation is indicated by the dotted line on the concentration-response curve. Scale bar = 40µm for all images. White arrows indicate nuclear signal localisation.

Both TGFβ and GDF-15 induced similar secretory profiles in brain pericytes (Figure 3). The concentrations of MCP-1 and IL-6 increased over time with both TGFβ and GDF-15 treatment (Figure 3). No change was seen in the concentration of RANTES, sICAM, sVCAM, TNF, IL-8, or VEGF (Figure 3).

**Figure 3:**
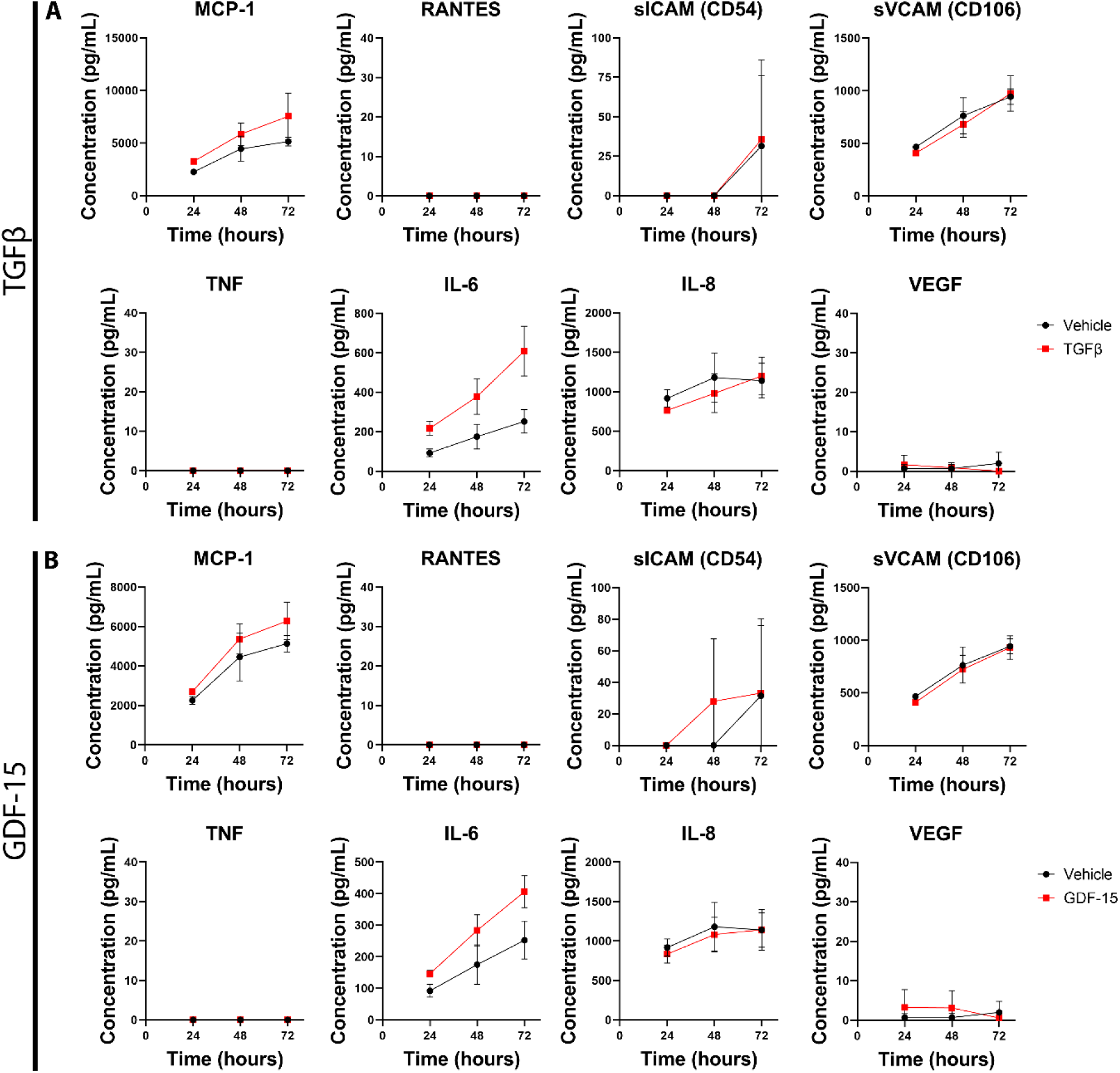
TGFβ cytokine superfamily cytokines TGFβ and GDF-15 stimulate inflammatory secretions in brain pericytes. Graphs demonstrating the concentration of MCP-1, RANTES, sICAM, sVCAM, TNF, IL-6, IL-8, and VEGF in response to TGFβ treatment (A, 200pM or 5ng/mL) or GDF-15 treatment (B, 8.13nM or 200ng.mL) over 72 hours. The presented data is collected from two experimental repeats.

### Inflammatory factors IL-17, IL-27, and IFNγ induce changes in primary pericyte inflammatory phenotype

Primary pericytes demonstrated responses to the inflammatory cytokines IL-17, IL-27, and IFNγ (Figures 4-5). No transcription factor translocation was observed with IL-17 treatment (Figure 4A/B). IL-27 induced nuclear localisation of STAT1 in 35.3% of primary pericytes with an EC50 of 82.0pM, a partial response comparable to that of IL-1β or TNF treatment observed by our lab previously (14) (Figure 4C/D). A small proportion of pericytes exhibited a C/EBP-δ response to IL-27 treatment with an EC50 of 27.6pM (Figure 4C/D). While only 14.1% of pericytes responded to IL-27 treatment, this is likely an underestimation of the true number of responsive cells due to the nuclear C/EBP-δ present in 70% of primary pericytes in vehicle-treated samples (Figure 4C/D). Interestingly, this could suggest that a component of the Sciencell pericyte medium induces C/EBP-δ signalling. Previous studies have found IFNγ treatment to exhibit a STAT1 response in primary human brain pericytes (14). Here, we find this signalling pathway is conserved in foetal human brain pericytes. STAT1 translocation was strongly induced by IFNγ, with translocation demonstrated in 89.0% of cells with an average EC50 of 171pM (Figure 4E-F). No C/EBP-δ response was observed with IFNγ treatment.

**Figure 4:**
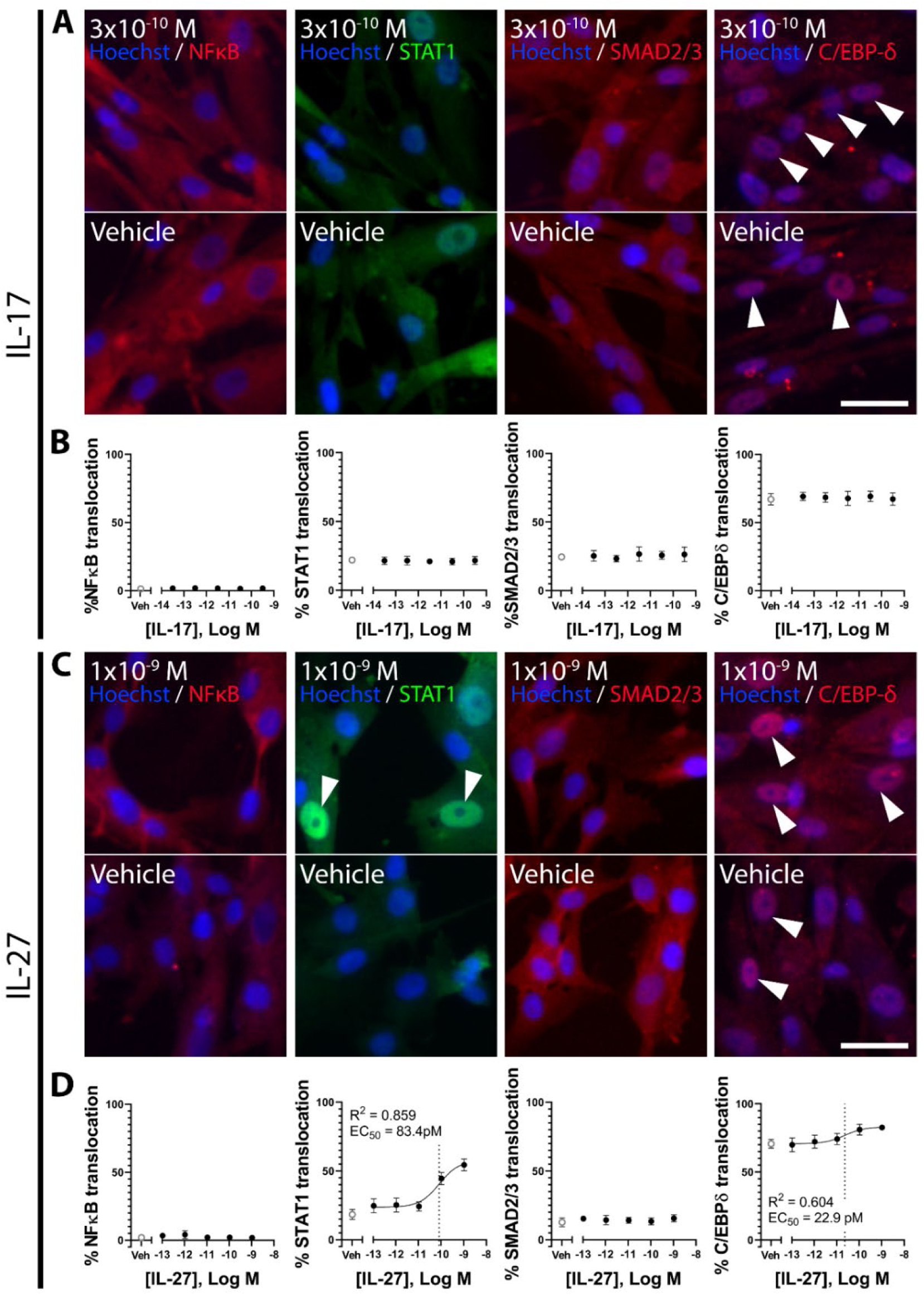
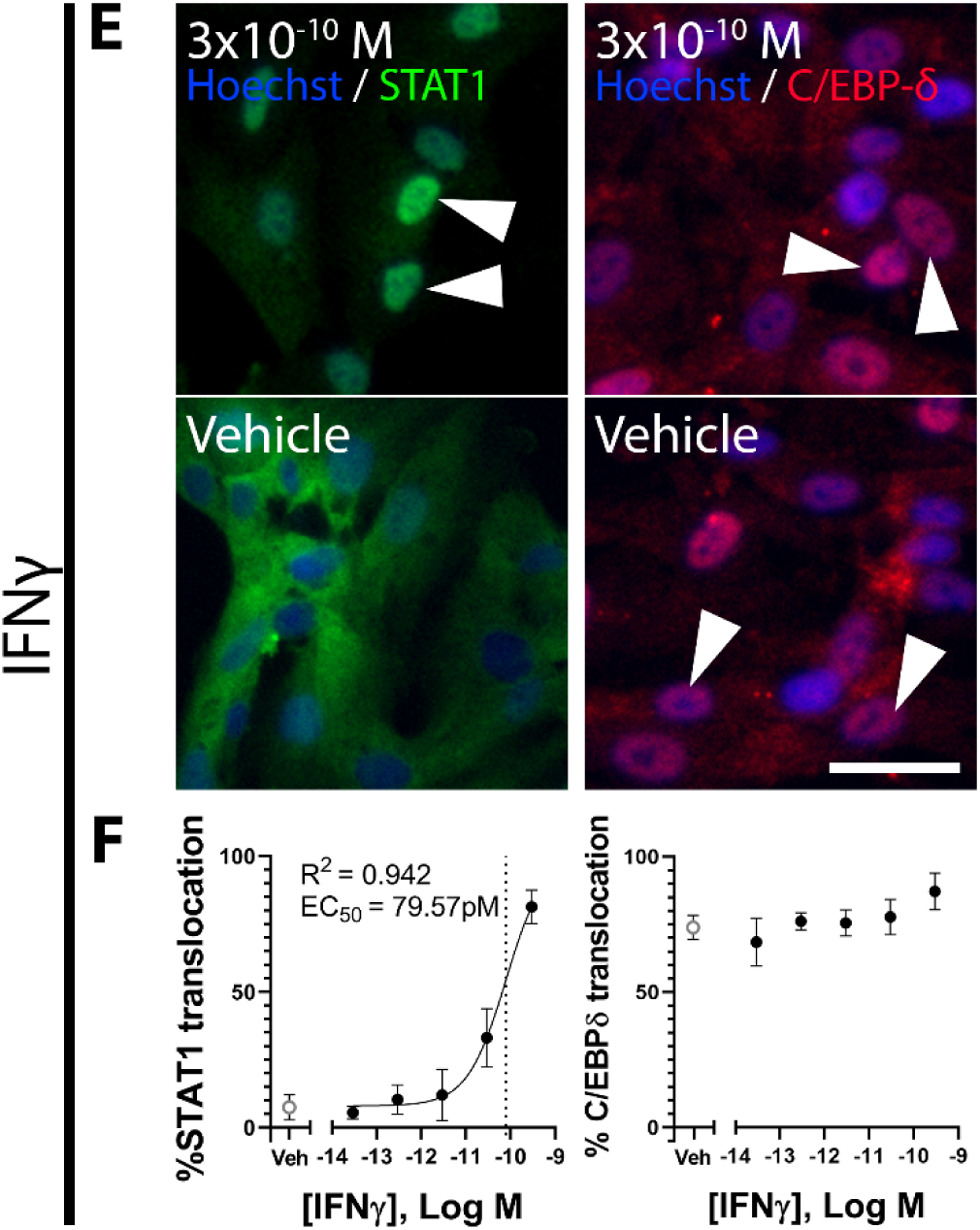
Signalling responses to treatment with IL-17, IL-27 and IFNγ. Immunofluorescence images demonstrating NFκB, STAT1, SMAD2/3, and C/EBP-δ localisation in response to IL-17 (A), IL-27 (C) or IFNγ (E) in primary brain pericytes. Images are quantified using MetaXpress to generate concentration-response curves (B,D,F). Data presented is one representative experiment of at least two experimental repeats for each cytokine treatment (see supplemental table 4). The EC_50_ for nuclear localisation is indicated by the dotted line on the concentration-response curve. Scale bar = 40µm for all images. White arrows indicate nuclear signal localisation.

**Figure 5:**
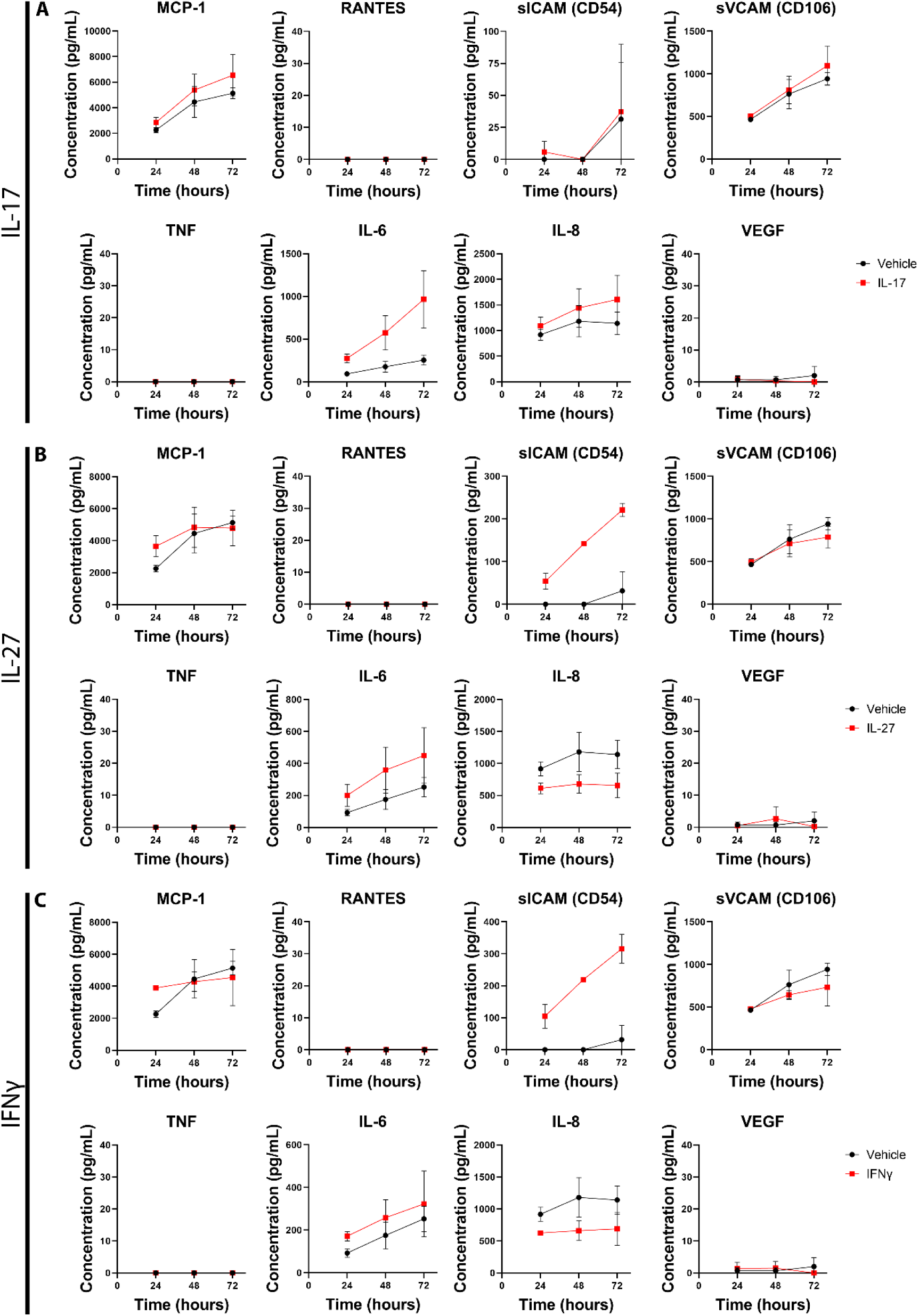
IL-17, IL-27, and IFNγ induce secretory changes in brain pericytes. Graphs demonstrating the concentration of MCP-1, RANTES, sICAM, sVCAM, TNF, IL-6, IL-8, and VEGF in response to IL-17 treatment (A, 316pM or 5ng/mL), IL-27 treatment (B, 1nM or 50ng/mL), or IFNγ treatment (C, 296pM or 5ng/mL) over 72 hours. The presented data is collected from two experimental repeats.

Following investigation of transcription factor activation, IL-17, IL-27, and IFNγ-mediated pericyte secretions were assessed. Interestingly, although no transcription factor activation was observed with IL-17 treatment, changes in the concentration of local inflammatory factors was observed. IL-17 induced an increase in MCP-1, IL-6, and IL-8 concentrations across 72 hours, indicative of a proinflammatory response (Figure 5A). No change was observed in the concentrations of RANTES, ICAM, VCAM, TNF, or VEGF (Figure 5A). IL-27 induced changes in the concentrations of ICAM, IL-6, and IL-8 (Figure 5B). Increases in the concentration of ICAM and IL-6 were observed across the 72 hours with IL-27 treatment (Figure 5B). Interestingly, the concentration of IL-8 was reduced with IL-27 treatment across the 72 hours (Figure 5B). IFNγ treatment exhibited a similar reduction in IL-8, indicating that this reduction may be a result of STAT1 translocation. Treatment with IFNγ also increased observed concentrations of sICAM, and IL-6 (Figure 5C). No change was observed for MCP-1, RANTES, sVCAM-1, TNF, or VEGF. These results show activity of inflammatory cytokines on primary brain pericytes, but also show anti-chemotactic activity of IL-27 and IFNγ through reduction on IL-8 secretion.

### IFNγ and IL-27 reduce phagocytosis in primary pericytes

Brain pericytes are capable of phagocytosis which could contribute to the hiding of tumour antigen through increased removal of tumour debris (16, 18). We investigated the effect of brain cancer-derived factors on pericyte phagocytosis. Fluorescent beads were incubated with primary pericytes *in vitro*. Phagocytic cells exhibited intracellular fluorescence which correlated with the rate of phagocytosis when investigated by flow cytometry and could be manipulated in a concentration-dependent manner (Figure S16). 83.7% of vehicle-treated primary pericytes exhibited phagocytic activity (Figure 6). There was no change in pericyte phagocytic activity following treatment with TGFβ, GDF-15, IL-17, fractalkine, MIF, PDGF-aa, CHI3L1, osteopontin, IGFBP2, or IGFBP3 (Figure 6A-C, Figure S17).

**Figure 6:**
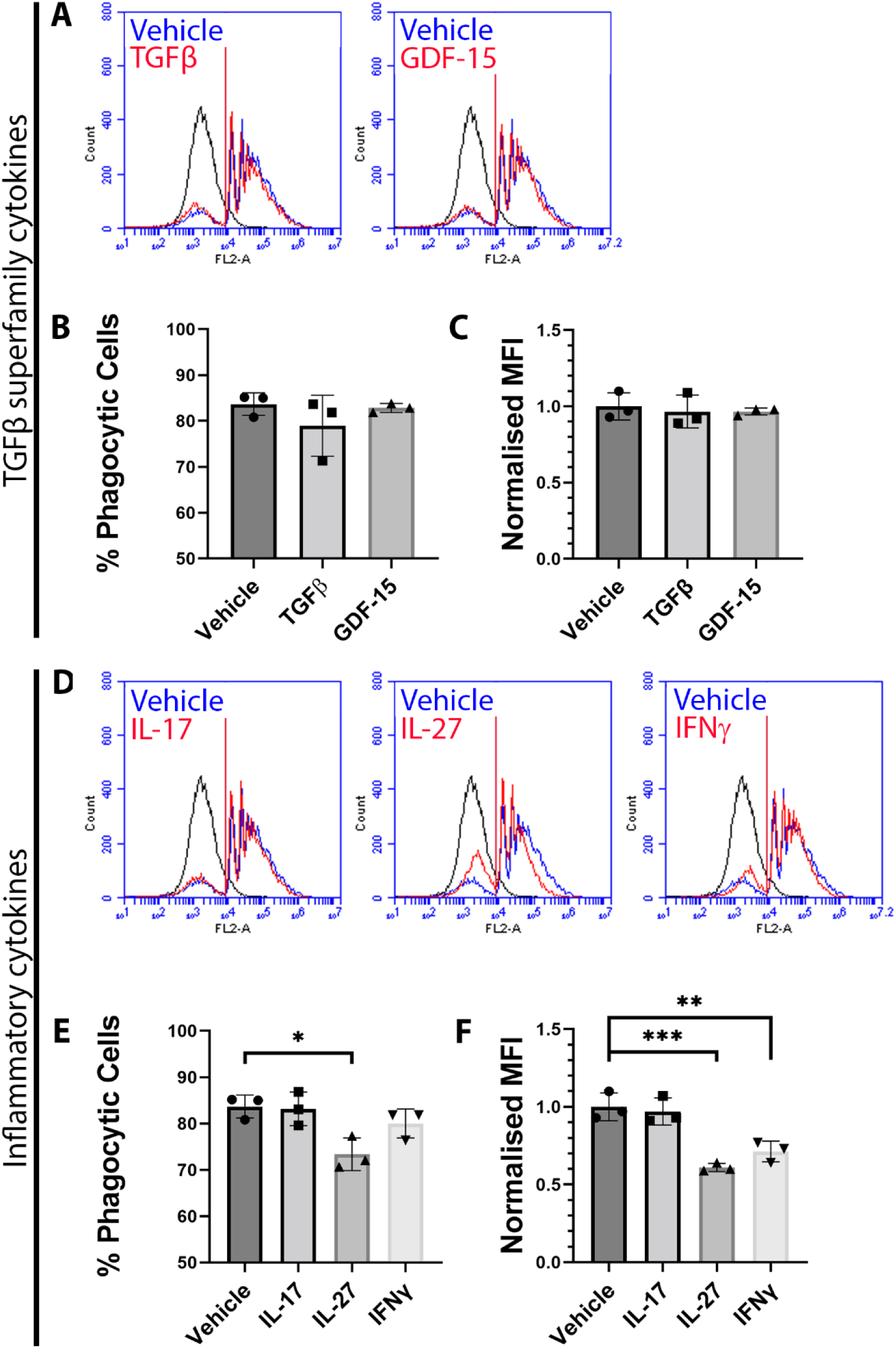
IFNγ and IL-27 treatment reduced phagocytosis in primary brain pericytes. Flow cytometry histo-plots showing the fluorescence of treated (red) and vehicle (blue) cell populations after incubation with fluorescent beads for 24 hours (A,D). Cellular auto-fluorescence can be seen in black. The vertical red line indicates the fluorescent threshold for cells to be considered phagocytic. Quantification of the histo-plots shows the percentage of phagocytic cells (B,E) and the normalised mean fluorescent intensity (MFI) of phagocytic cells (C,F). The presented data is collected from three experimental repeats. The significance of each treatment compared to vehicle was determined using a one-way ANOVA with multiple comparisons. *: p<0.05, **: p<0.01, ***: p<0.001.

A reduction in pericyte phagocytosis was exhibited with IL-27 and IFNγ treatments. IL-27 treatment induced the largest decrease, reducing the proportion of phagocytic cells from 83.7% to 73.4%, with phagocytic cells exhibiting a normalised MFI of 0.61 (Figure 6D-F). IFNγ treatment reduced the proportion of phagocytic cells from 87.3% to 80.0%, with phagocytic cells exhibiting a normalised MFI of 0.71 (Figure 6D-F). These data show that pericyte phagocytosis is resistant to many inflammatory stimuli, but can be reduced with the IL-27 or IFNγ treatment.

## Discussion

Brain pericytes occupy a critical vascular niche which could greatly help or hinder brain tumour progression. However, little is currently known about how brain pericytes react to the tumour microenvironment. We assessed the effect of individual cancer-derived factors on brain pericyte function. We found that TGFβ and GDF-15 inhibited C/EBP-δ signalling, offering a mechanism behind the TGFβ-induced secretion of IL-6 seen in brain pericytes, discussed below. The STAT1 activating cytokines IFNγ and IL-27 strongly reduced phagocytosis, indicating signalling pathways which may reduce phagocytosis in brain pericytes. Finally, the majority of tumour-derived factors did not produce a measurable change to brain pericyte activity, suggesting that these factors may act on other cell types such as tumour-associated macrophages (TAMs), or require co-stimulation with other factors commonly found within the tumour milieu. Together, these results offer new insight into foundational brain pericyte biology and indicate potential roles for pericytes in brain tumour progression.

TGFβ has been well studied in pericyte biology due to its role in pericyte differentiation, producing a smooth-muscle-like phenotype, as well as a pleiotropic but overall anti-inflammatory role during inflammation (16, 17). Our study finds that TGFβ inhibits C/EBP-δ, which may be causative of TGFβ-induced secretion of IL-6 and IL-8 (15, 16). C/EBP-δ exhibits anti-inflammatory effects in brain pericytes, therefore its inhibition potentiates the expression and secretion of numerous inflammatory chemokines and adhesion molecules such as sICAM-1, MCP-1, IL-6, and IL-8 (15). This reveals a potential mechanism through which TGFβ can act synergistically with IL-1β or other inflammatory stimuli to promote the secretion of chemotactic molecules and drive inflammation. In the context of GBM, increased expression of TGFβ is correlated with maintenance of the GBM stem cell phenotype and immunosuppressive microenvironment, as well as increased invasiveness, poor patient prognosis, and reduced overall survival (19–22). TGFβ activity in brain pericytes may play a role in tumour invasiveness through regulation of matrix metalloproteinase-2 (MMP-2) expression (15, 23). Indeed, increased MMP-2 expression has been demonstrated in human brain pericytes with C/EBP-δ inhibition (15). This reveals C/EBP-δ as a potential target to reduce GBM tumour invasiveness.

The TGFβ-induced inhibition of C/EBP-δ translocation would not be evident if the primary pericytes didn’t exhibit such a high proportion of C/EBP-δ-positive cells in vehicle-treated samples. While nuclear C/EBP-δ did lead to this serendipitous finding, it posed a significant limitation when investigating C/EBP-δ induction from other factors. Nuclear translocation of C/EBP-δ was induced by IFNγ and IL-27 in a subset of primary pericytes, however, the proportion of C/EBP-δ-responsive cells is unknown. Additionally, C/EBP-δ translocation in a small subpopulation could be completely masked by the high proportion of C/EBP-δ-positive nuclei basally. We predict that the nuclear C/EBP-δ is a result of the influx of fresh Sciencell medium with treatment, given that the proportion of C/EBP-δ-positive nuclei increases between 1 and 4 hours of treatment (Figure S1). Sciencell pericyte medium uses a proprietary pericyte growth supplement containing growth factors, hormones, and proteins likely responsible for C/EBP-δ translocation. Furthermore, it is possible that trophic factors within the Sciencell medium are responsible for the relatively high concentrations of IL-8 and sVCAM seen in vehicle-treated medium of brain pericytes. Again, this lead to the serendipitous finding that IFNγ and IL-27 reduce the secretion of these cytokines in brain pericytes. Given the common use of Sciencell human brain pericytes, further work must be invested to uncover the role of C/EBP-δ in pericytes at rest and whether this recapitulates the *in vivo* pericyte phenotype.

We also found TGFβ-like responses from GDF-15 including SMAD2/3 translocation as well as C/EBP-δ inhibition. This adds to the conflicting evidence regarding GDF-15 activity, as studies have shown TGFβ contamination in commercially available GDF-15 protein (24). Peprotech, who supplied the GDF-15, presented data which showed a lack of TGFβ-like bioactivity, as well as known GDF-15 activity from the GDF-15 lot used in this study (Figures S18-S19). This corroborates previous findings that GDF-15 induces SMAD2 phosphorylation through both ALK and ERK1/2 signalling (25). Importantly, this signalling may create an autocrine/paracrine loop whereby GDF-15 signalling promotes further GDF-15 secretion (25). Brain pericytes may contribute to GDF-15 secretion following initial exposure to GDF-15, promoting the immune-suppressive phenotype present in GBM, while also likely promoting cancer stem cell-like properties mediated by FOXM1 (25, 26). These results indicate a novel pathway through which brain pericytes may support brain tumour progression.

GBM typically overexpress chemoattractants such as fractalkine, MCP-1, and IL-8 to recruit tumour-associated macrophages (TAMs), whilst simultaneously expressing proteins such as CD47 to reduce cell phagocytosis, thereby preventing TAMs from processing and presenting antigens (27, 28). As brain pericytes do not express the cellular machinery for canonical antigen presentation, brain pericytes may phagocytose tumour antigens to instead sequester it, preventing their acquisition by antigen presenting cells, thereby acting as a source of immunosuppression. Primary pericytes demonstrated significant inhibition of phagocytosis with IFNγ and IL-27 treatment. As STAT1 has been shown to reduce phagocytosis in immune cells such as macrophages and dendritic cells, as well as other epithelial cell types, this is likely also regulating phagocytosis in IFNγ- and IL-27-treated brain pericytes (29–31). Inhibition of phagocytosis could prevent brain pericytes from hiding tumour antigen, thereby allowing for canonical antigen processing of tumour antigens in brain cancers. However, overall STAT1 demonstrates pro-tumourigenic effects and is often upregulated in glioblastoma (32, 33). In fact, IGFBP3 induces STAT1 expression in glioblastoma cells (32). Phagocytosis is significantly reduced in IL-27-treated pericytes, suggesting additional mechanisms by which this cytokine induces a loss in phagocytosis. Understanding these mechanisms may reveal alternative targets to reduce stromal phagocytosis, thereby alleviating some immunosuppressive functions of glioblastoma.

Of note, most of the assessed cancer-associated factors did not induce nuclear translocation of major signalling factors in brain pericytes. This could be due to the acute timepoint (1 hour) used to assess NFκB, STAT1, and SMAD2/3 signalling, which may not capture the full response of some signalling pathways. However, this acute timeframe is consistent with direct receptor-mediated signalling, indicating perhaps that these factors may potentiate or inhibit some signalling pathways instead of activating them directly. For example, CHI3L1 has been found to sensitise IL-13–mediated inflammatory responses, reducing inflammatory secreted products (34, 35). Additionally, IL-17 is a potent inflammatory activator *in vivo*, however demonstrates only mild activity *in vitro*, suggesting that other factors are required for IL-17 to function optimally (36). While it is important to investigate the action of cancer-associated factors individually, this doesn’t recapitulate the complexity of the GBM microenvironment. Given that these factors didn’t activate major signalling pathways on their own, this highlights the need for future experiments to use multiple treatments simultaneously to investigate synergistic or inhibitory effects of these factors.

## Conclusions

The work presented here finds novel signalling responses in primary pericytes when stimulated with cancer-associated and inflammatory factors. We found TGFβ to inhibit the nuclear translocation of C/EBP-δ, revealing a candidate mechanism for the potentiation of brain pericyte inflammation. GDF-15 shared TGFβ signalling pathways, indicating that it may support tumour progression through similar mechanisms. Finally, IL-27 generated a potent STAT1 response in brain pericytes, significantly reducing phagocytosis in these cells. These data show potential mechanisms by which brain pericytes contribute to aspects of inflammation and starts to uncover the supportive role brain pericytes may play in brain cancers.

## Ethics approval and consent to participate

not applicable.

## Availability of data and materials

The datasets used and/or analysed during the current study are available from the corresponding author on reasonable request.

## Author Contributions

<u>Conceptualization</u>, Samuel McCullough and E. Scott Graham; <u>Data</u> <u>curation</u>, Samuel McCullough; <u>Formal analysis</u>, Samuel McCullough, Eliene Albers, Bronwen Connor, and E. Scott Graham; <u>Funding acquisition</u>, Samuel McCullough and E. Scott Graham; <u>Investigation</u>, Samuel McCullough and Akshata Anchan; <u>Methodology</u>, Samuel McCullough, Akshata Anchan, Jane Yu and E. Scott Graham; <u>Project administration</u>, Eliene Albers, Bronwen Connor and E. Scott Graham; <u>Resources</u>, Samuel McCullough, Eliene Albers, Bronwen Connor and E. Scott Graham; <u>Software</u>, Akshata Anchan, Jane Yu and E. Scott Graham; <u>Supervision</u>, Eliene Albers, Bronwen Connor and E. Scott Graham; <u>Validation</u>, Samuel McCullough, Akshata Anchan, Jane Yu and E. Scott Graham; <u>Visualization</u>, Samuel McCullough; <u>Writing – original draft</u>, Samuel McCullough; <u>Writing – review & editing</u>, Samuel McCullough, Eliene Albers, Akshata Anchan, Bronwen Connor and E. Scott Graham. All authors have read and agreed to the published version of the manuscript.

## Acknowledgements

This work was funded by the Department of Anatomy and Medical Imaging, Faculty of Medical and Health Sciences, University of Auckland, New Zealand. Diagrams were generated using Biorender.

## List of abbreviations

BBB: Blood-brain barrier
CBA: Cytometric bead array
C/EBP-δ: CCAAT-enhancer-binding protein-δ
CHI3L1: Chitinase-3-like protein 1
DMSO: Dimethyl sulfoxide
EDTA: Ethylenediaminetetraacetic acid
FBS: Foetal bovine serum
GBM: Glioblastoma multiforme
GDF-15: Growth-differentiation factor-15
ICAM-1: Intercellular adhesion molecule 1
IFNγ: Interferon γ
IGFBP2: Insulin-like growth factor binding protein-2
IGFBP3: Insulin-like growth factor binding protein-3
IL-1β: Interleukin-1β
IL-6: Interleukin-6
IL-8: Interleukin-8
IL-17: Interleukin-17
IL-27: Interleukin-27
MCP-1: Monocyte chemotactic protein-1
MFI: Mean fluorescent intensity
MIF: Macrophage migration inhibitory factor
NFκB: Nuclear factor kappa-light-chain-enhancer of activated B cells
PBS: Phosphate buffered saline
PDGF-aa: Platelet-derived growth factor-aa
RANTES: Regulated upon Activation, Normal T Cell Expressed and Presumably Secreted
SMAD2/3: Mothers against decapentaplegic homolog 2/3
STAT1: Signal transducer and activator of transcription 1
STAT3: Signal transducer and activator of transcription 3
TAM: Tumour-associated macrophages
TGFβ: Transforming growth factor β
TNF: Tumour necrosis factor
VCAM-1: Vascular cell adhesion molecule 1
VEGF: Vascular endothelial growth factor

