## Supplementary material for "Screening activity of brain cancer-derived factors on primary human brain pericytes"

### Supplemental material

**Supplemental table 1: Summary of inflammatory treatments used in this study.**

| Treatment | ICC max conc. (M) | ICC max conc. (ng/mL) | ICC treatment time | CBA and phagocytosis treatment conc. (M) | Supplier | Cat. # |
|---|---|---|---|---|---|---|
| IL-1β | $2.89 \times 10^{-10}$ | 5 | 1h, 4h | $2.89 \times 10^{-10}$ | Peprotech | 200-01B |
| TNF | $2.89 \times 10^{-10}$ | 5 | 1h, 4h | $2.89 \times 10^{-11}$ | Peprotech | 300-01A |
| IFNγ | $2.96 \times 10^{-10}$ | 5 | 1h, 4h | $2.96 \times 10^{-10}$ | R&D Systems | 285-IF |
| TGFβ | $2.00 \times 10^{-10}$ | 5 | 1h, 4h | $2.00 \times 10^{-10}$ | Peprotech | 100-21 |
| CHI3L1 | $1.74 \times 10^{-9}$ | 80 | 1h, 4h | $1.74 \times 10^{-9}$ | Abcam | ab140057 |
| GDF-15 | $8.13 \times 10^{-9}$ | 200 | 1h, 4h | $8.13 \times 10^{-9}$ | Peprotech | 120-28C |
| Fractalkine | $5.88 \times 10^{-10}$ | 5 | 1h, 4h | $5.88 \times 10^{-10}$ | Peprotech | 300-31 |
| MIF | $3.33 \times 10^{-10}$ | 5 | 1h, 4h | $3.33 \times 10^{-10}$ | Peprotech | 300-69 |
| IL-17A | $3.16 \times 10^{-10}$ | 5 | 1h, 4h | $3.16 \times 10^{-10}$ | Peprotech | 200-17 |
| IL-27 | $1.05 \times 10^{-9}$ | 50 | 1h, 4h | $1.05 \times 10^{-9}$ | Peprotech | 200-38 |
| PDGF-aa | $1.75 \times 10^{-10}$ | 5 | 1h, 4h | $1.75 \times 10^{-10}$ | Peprotech | 100-13A |
| Osteopontin | $2.97 \times 10^{-9}$ | 100 | 1h, 4h | $2.97 \times 10^{-9}$ | Peprotech | 120-35 |
| IGFBP2 | $1.59 \times 10^{-9}$ | 50 | 1h, 4h | $1.59 \times 10^{-9}$ | Peprotech | 350-06B |
| IGFBP3 | $3.47 \times 10^{-9}$ | 100 | 1h, 4h | $3.47 \times 10^{-9}$ | Peprotech | 100-08 |

Immunocytochemistry treatment time was 1 hour (1h) for investigation of NFκB, STAT1, and SMAD2/3, and 4 hours (4h) for investigation of C/EBP-δ.

**Supplemental table 2: List of immunocytochemistry dilutions used in this study.**

| Stain | Dilution | Catalogue # | Company | Secondary Ab. | Secondary Dilution |
|---|---|---|---|---|---|
| Hoechst | 1:10,000 | H3570 | Invitrogen | n/a | n/a |
| NFκB | 1:500 | sc-8008 | Santa Cruz Biotechnology | 594 Donkey α Mouse | 1:500 |
| STAT1 | 1:500 | 14994 | Cell Signalling Technology | 488 Donkey α Rabbit | 1:500 |
| SMAD2/3 | 1:500 | sc-133098 | Santa Cruz Biotechnology | 594 Donkey α Mouse | 1:500 |
| C/EBP-δ | 1:250 | sc-365546 | Santa Cruz Biotechnology | 594 Donkey α Mouse | 1:500 |

**Supplemental table 3: List of CBA kits used in this study.**

| CBA Kit Target | Catalogue # | Bead position |
|---|---|---|
| MCP-1 | 558287 | D8 |
| RANTES | 558324 | D4 |
| sICAM-1 | 560269 | A4 |
| sVCAM-1 | 560427 | D6 |
| TNF | 558273 | D9 |
| IL-6 | 558276 | A7 |
| IL-8 | 558277 | A9 |
| VEGF | 558336 | B8 |

**Supplemental table 4: Summary of transcription factor responses to treatment.**

| | NFκB | | | STAT1 | | | SMAD2/3 | | | C/EBP-δ | | |
|---|---|---|---|---|---|---|---|---|---|---|---|---|
| | EC50 (pM) | % Responsive cells | Experimental repeats | EC50 (pM) | % Responsive cells | Experimental repeats | EC50 (pM) | % Responsive cells | Experimental repeats | EC50 / IC50 (pM) | % Responsive cells | Experimental repeats |
| **IL-1β** | 1.75 ± 1.37 | 82.6 ± 12.4 | 4 | 3.57 ± 0.872 | 26.9 ± 2.59 | 3 | n/a | n/a | 3 | 3.92 ± 0.80 | 34.5 ± 8.2 | 2 |
| **TNF** | 4.9 ± 2.47 | 77.2 ± 6.83 | 3 | 0.924 ± 0.837 | 18.3 ± 2.06 | 3 | n/a | n/a | 2 | n/a | n/a | 2 |
| **IFNγ** | n/a | n/a | n/a | 171 ± 130 | 89.0 ± 5.73 | 2 | n/a | n/a | n/a | 85.6 ± 19.4 | 19.9 ± 3.18 | 2 |
| **TGFβ** | n/a | n/a | 2 | n/a | n/a | 2 | 12.1 ± 1.05 | 68.3 ± 16.8 | 2 | 15.3 ± 9.72* | 17.9 ± 1.34* | 2 |
| **GDF-15** | n/a | n/a | 3 | n/a | n/a | 3 | 1515 ± 1851 | 65.7 ± 29.8 | 3 | n/a # | n/a # | 2 |
| **IL-17** | n/a | n/a | 2 | n/a | n/a | 2 | n/a | n/a | 2 | n/a | n/a | 2 |
| **IL-27** | n/a | n/a | 3 | 82 ± 2.05 | 35.3 ± 1.91 | 4 | n/a | n/a | 4 | 27.6 ± 6.65 | 14.1 ± 2.33 | 2 |

Results are displayed as the average between the experimental repeats ± standard deviation. Cells marked with "n/a" did not exhibit a response or a response was not investigated. "*" indicates $IC_{50}$ instead of $EC_{50}$. "#" indicates that a response was observed but couldn't be quantified due to an incomplete concentration-response curve.

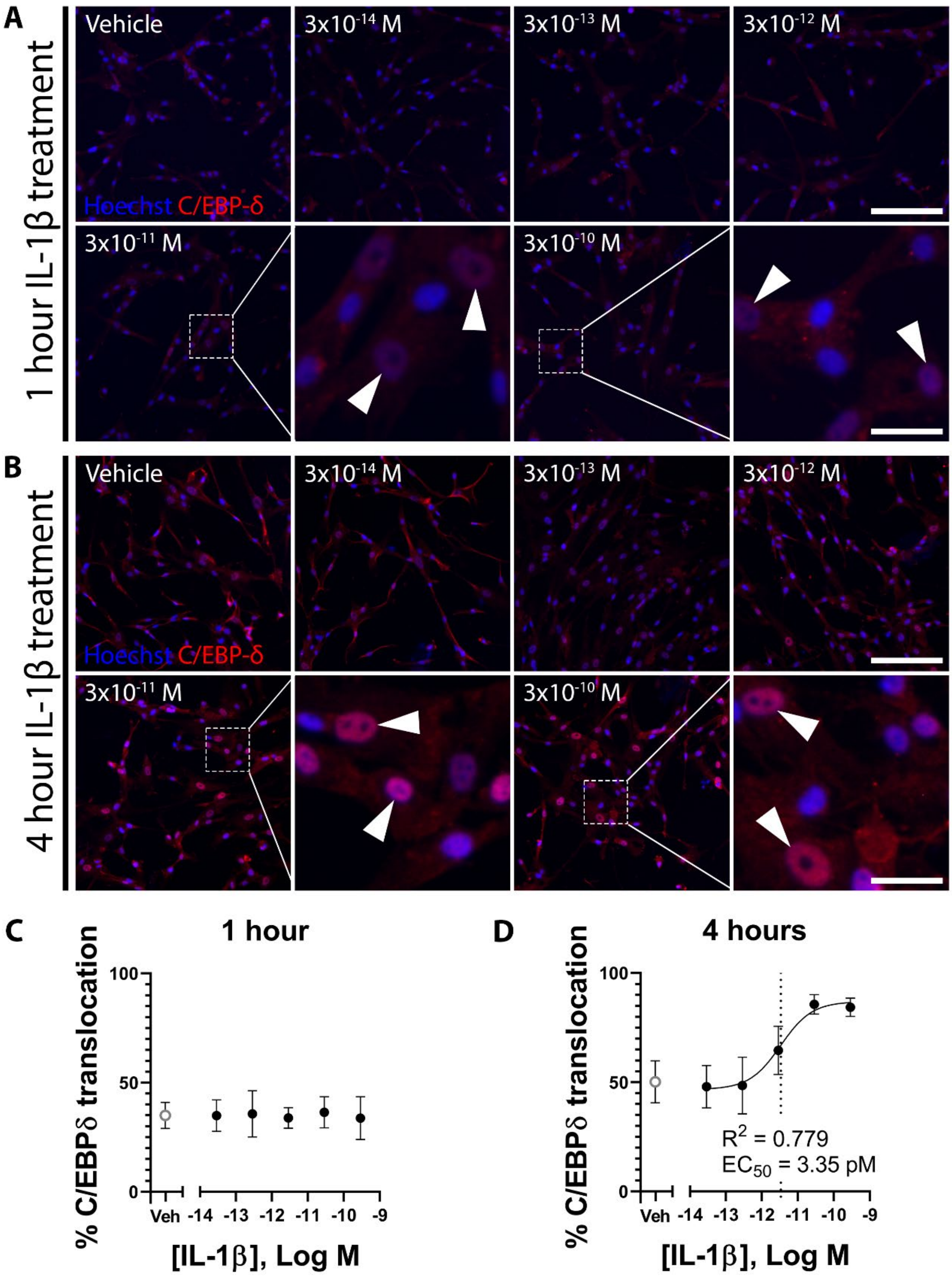


**Supplemental figure 1: C/EBP-δ translocates to the nucleus after four hours of IL-1β treatment in primary pericytes.** Immunofluorescence images demonstrating nuclear localisation of C/EBP-δ at one hour (A), or nuclear translocation after four hours (B) in response to increasing concentrations of IL-1β in primary pericytes. Arrows indicate examples of pericytes showing nuclear localisation of C/EBP-δ. Images are quantified using MetaXpress to generate concentration-response curves which demonstrates no concentration-dependent response at one hour (C), but an $EC_{50}$ of 3.35pM (D, dotted line) after four hours. Data presented is one representative experiment of two experimental repeats. Scale bar = 200μm for all non-magnified images. Magnified scale bar = 40μm for all magnified images.

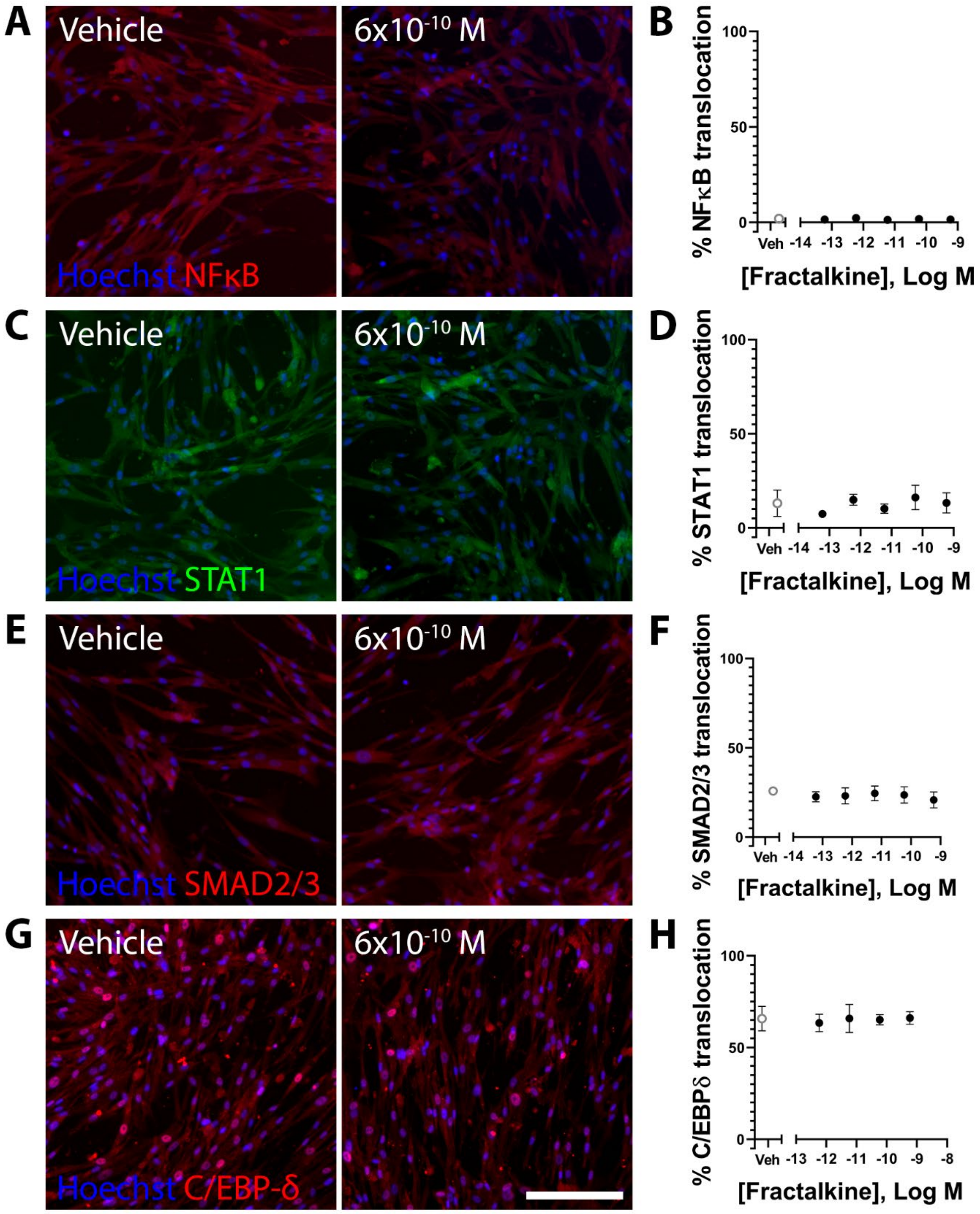


**Supplemental figure 2: Fractalkine treatment does not cause translocation of NFκB, STAT1, SMAD2/3 or C/EBP-δ in primary pericytes.** Immunofluorescence images demonstrating no change in NFκB (A,B), STAT1 (C,D), SMAD2/3 (E,F), or C/EBP-δ (G,H) localisation in response to increasing concentrations of fractalkine in primary pericytes (only highest concentration shown). Images are quantified using MetaXpress to generate concentration-response curves (B,D,F,H) which demonstrate no fractalkine-dependent change in signalling protein localisation. Data presented is one representative experiment of two experimental repeats. Scale bar = 200μm for all images.

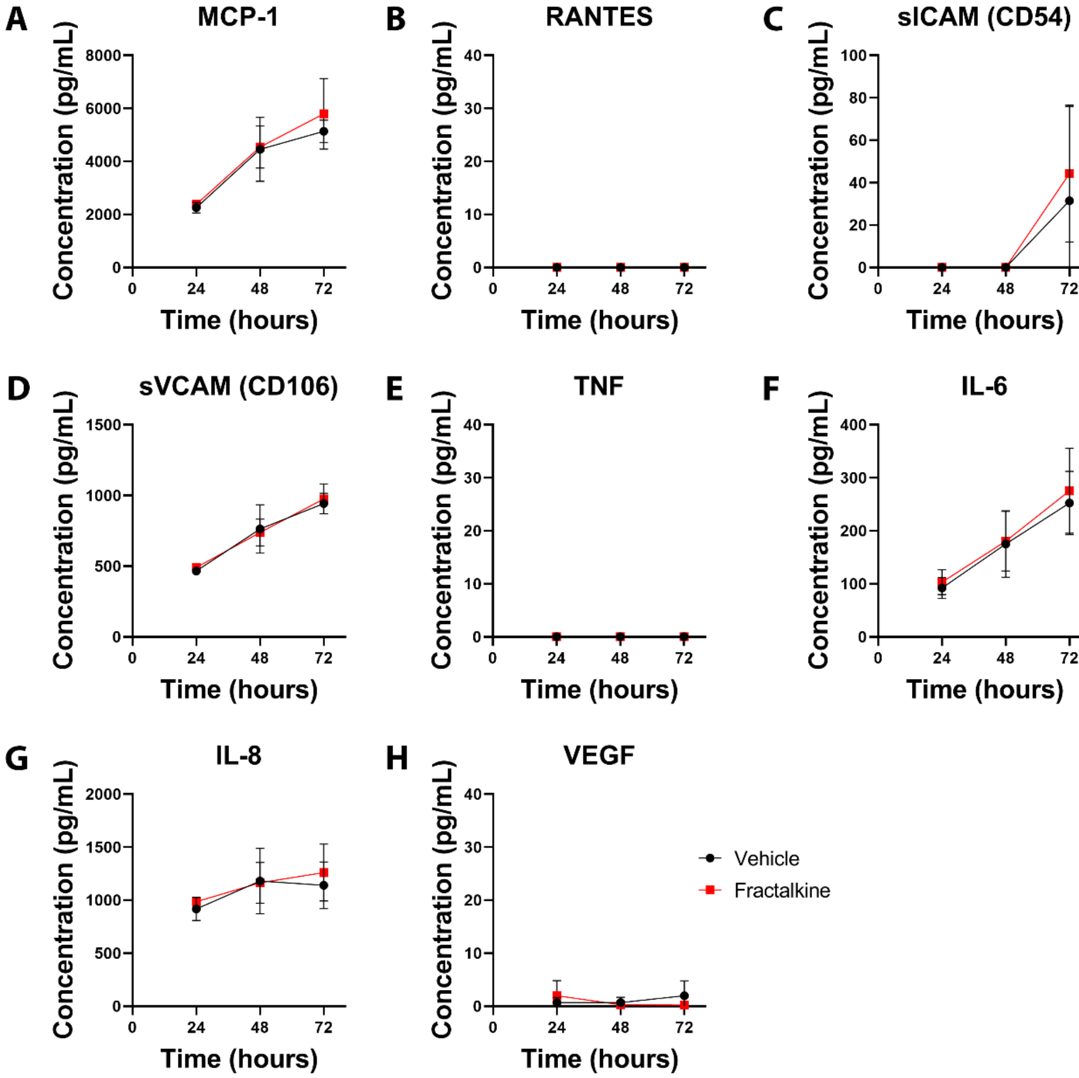


**Supplemental figure 3: Primary pericytes do not change secretion of inflammatory factors in response to fractalkine.** Pericyte medium was sampled with replacement 24, 48, and 72 hours after treatment with either fractalkine or vehicle. The concentration of MCP-1 (A), RANTES (B), sICAM (C), sVCAM (D), TNF (D), IL-6 (F), IL-8 (G), and VEGF (H) was investigated in each media sample by cytometric bead array. The presented data is collected from two experimental repeats.

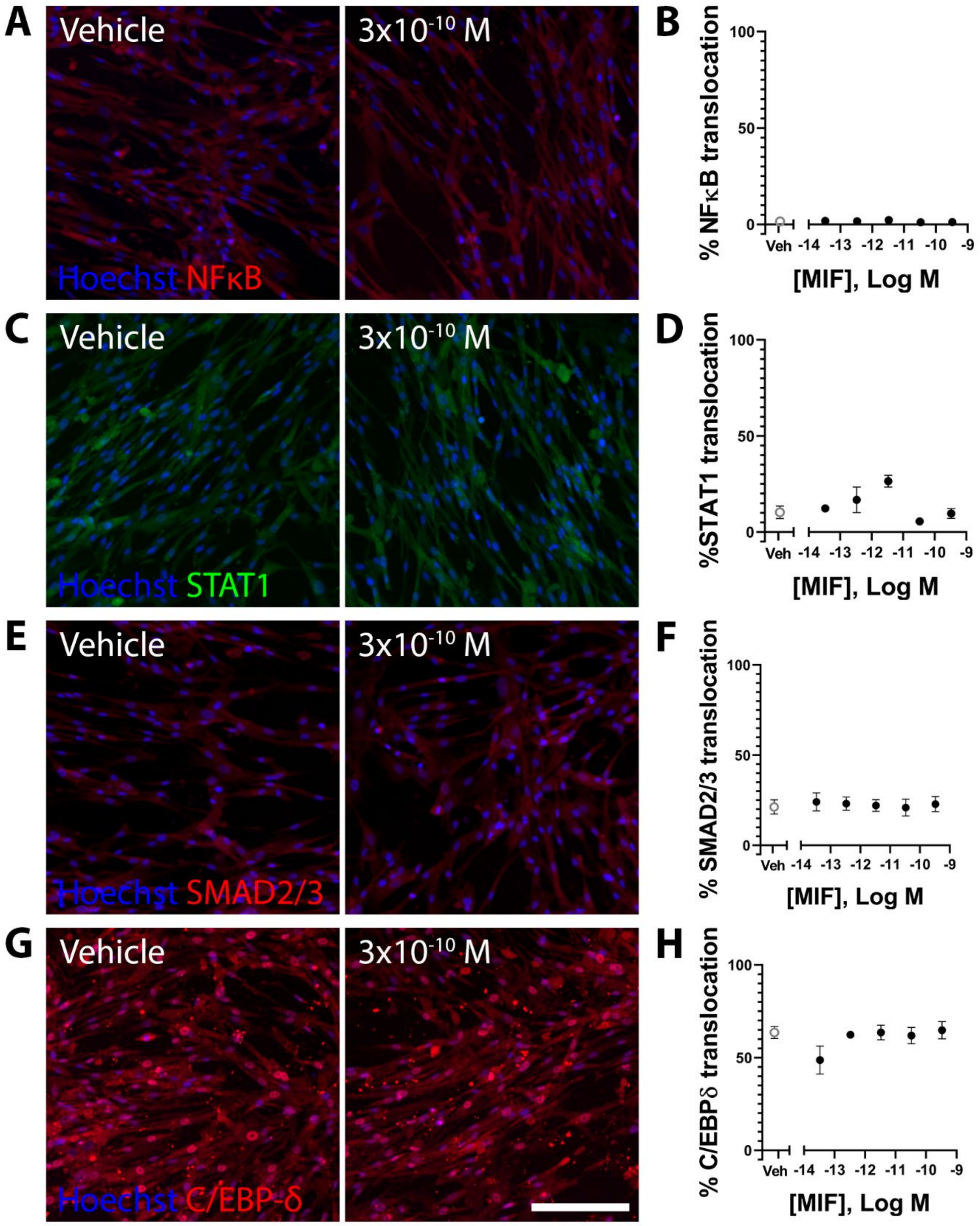


**Supplemental figure 4: MIF treatment does not cause translocation of NFκB, STAT1, SMAD2/3 or C/EBP-δ in primary pericytes.** Immunofluorescence images demonstrating no change in NFκB (A,B), STAT1 (C,D), SMAD2/3 (E,F), or C/EBP-δ (G,H) localisation in response to increasing concentrations of MIF in primary pericytes (only highest concentration shown). Images are quantified using MetaXpress to generate concentration-response curves (B,D,F,H) which demonstrate no MIF-dependent change in signalling protein localisation. Data presented is one representative experiment of two experimental repeats. Scale bar = 200μm for all images.

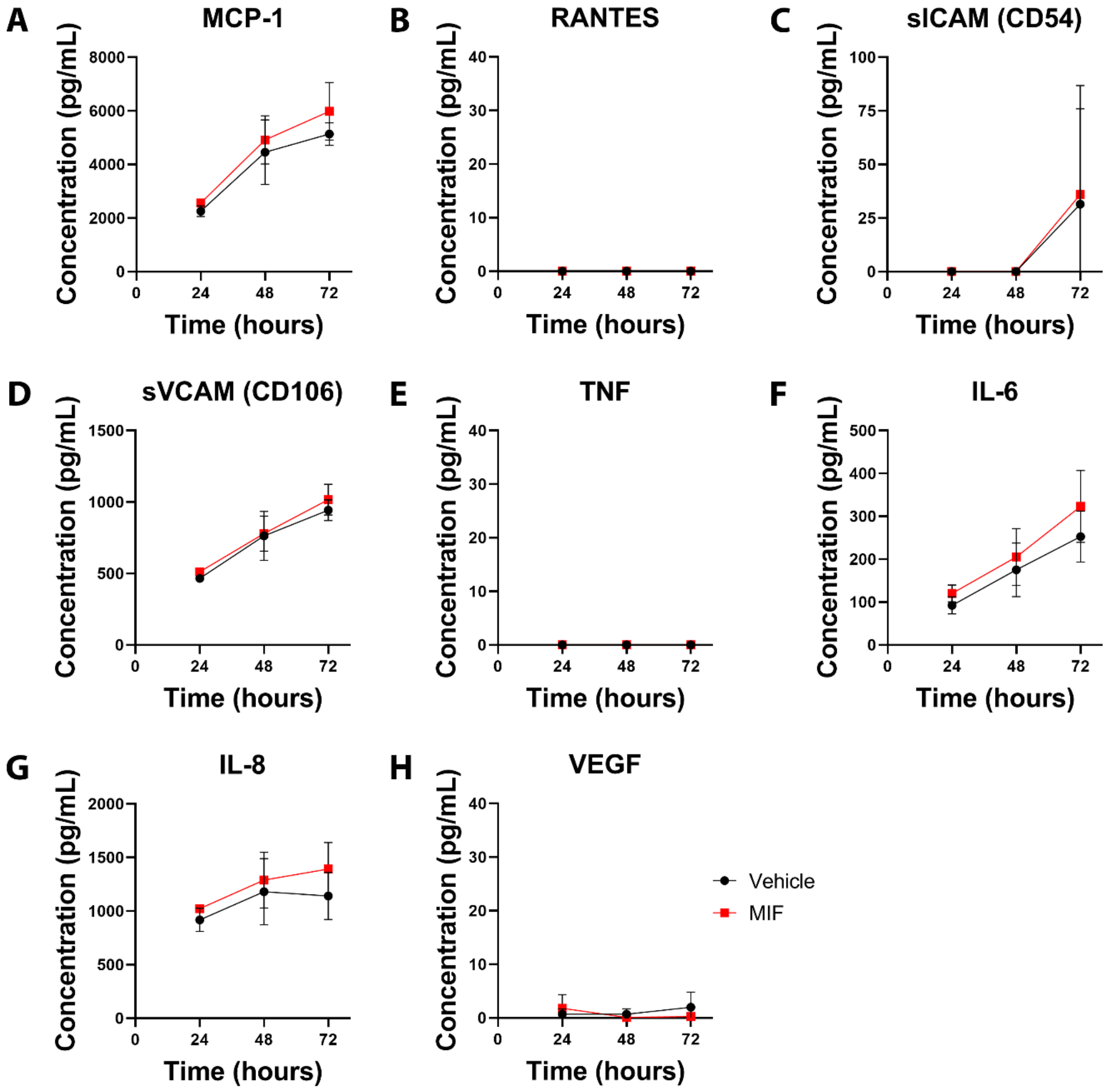


**Supplemental figure 5: Primary pericytes do not change secretion of inflammatory factors in response to MIF.** Pericyte medium was sampled with replacement 24, 48, and 72 hours after treatment with either MIF or vehicle. The concentration of MCP-1 (A), RANTES (B), sICAM (C), sVCAM (D), TNF (E), IL-6 (F), IL-8 (G), and VEGF (H) was investigated in each media sample by cytometric bead array. The presented data is collected from two experimental repeats.

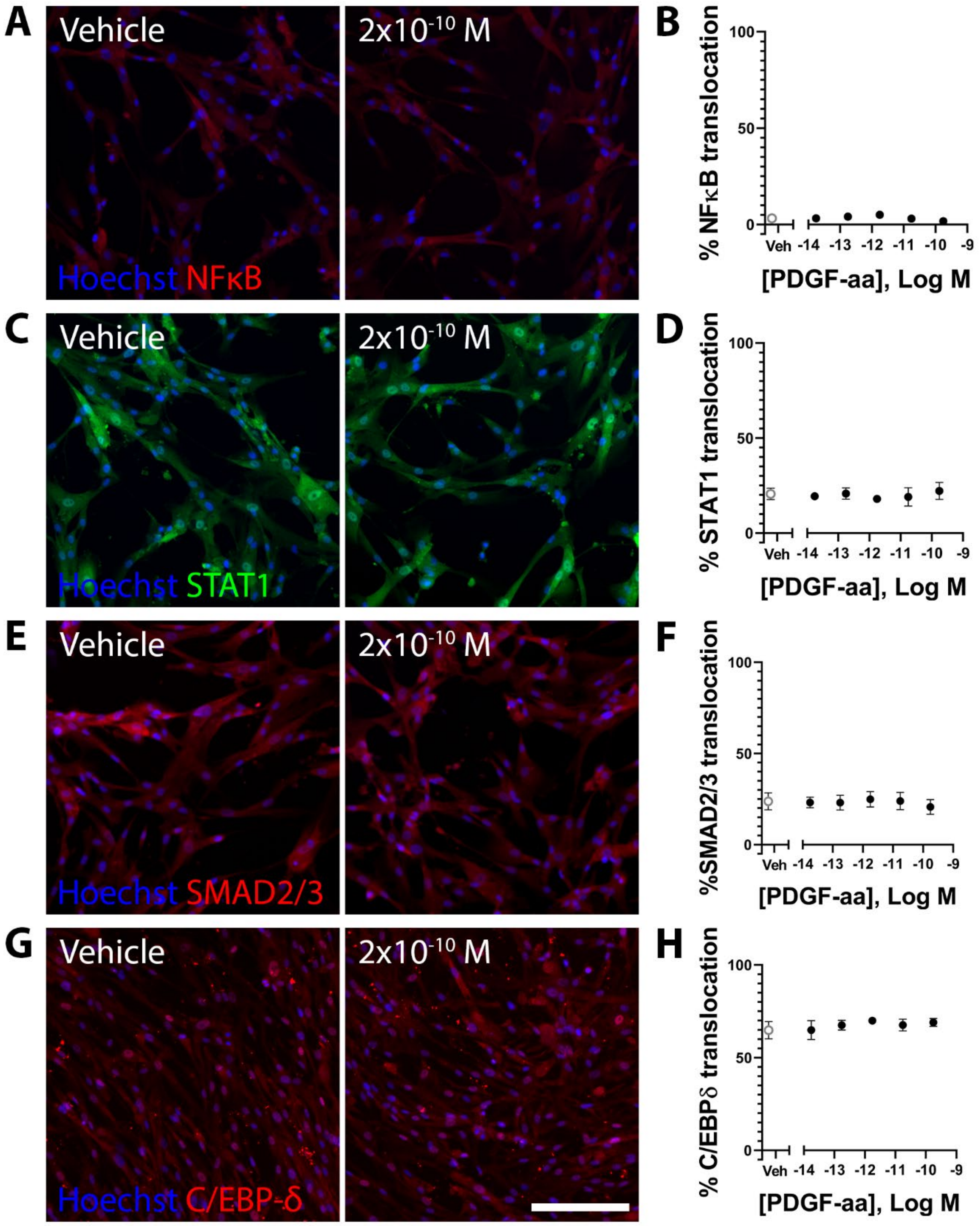


**Supplemental figure 6: PDGF-aa treatment does not cause translocation of NFκB, STAT1, SMAD2/3 or C/EBP-δ in primary pericytes.** Immunofluorescence images demonstrating no change in NFκB (A,B), STAT1 (C,D), SMAD2/3 (E,F), or C/EBP-δ (G,H) localisation in response to increasing concentrations of PDGF-aa in primary pericytes (only highest concentration shown). Images are quantified using MetaXpress to generate concentration-response curves (B,D,F,H) which demonstrate no PDGF-aa-dependent change in signalling protein localisation. Data presented is one representative experiment of two experimental repeats. Scale bar = 200μm for all images.

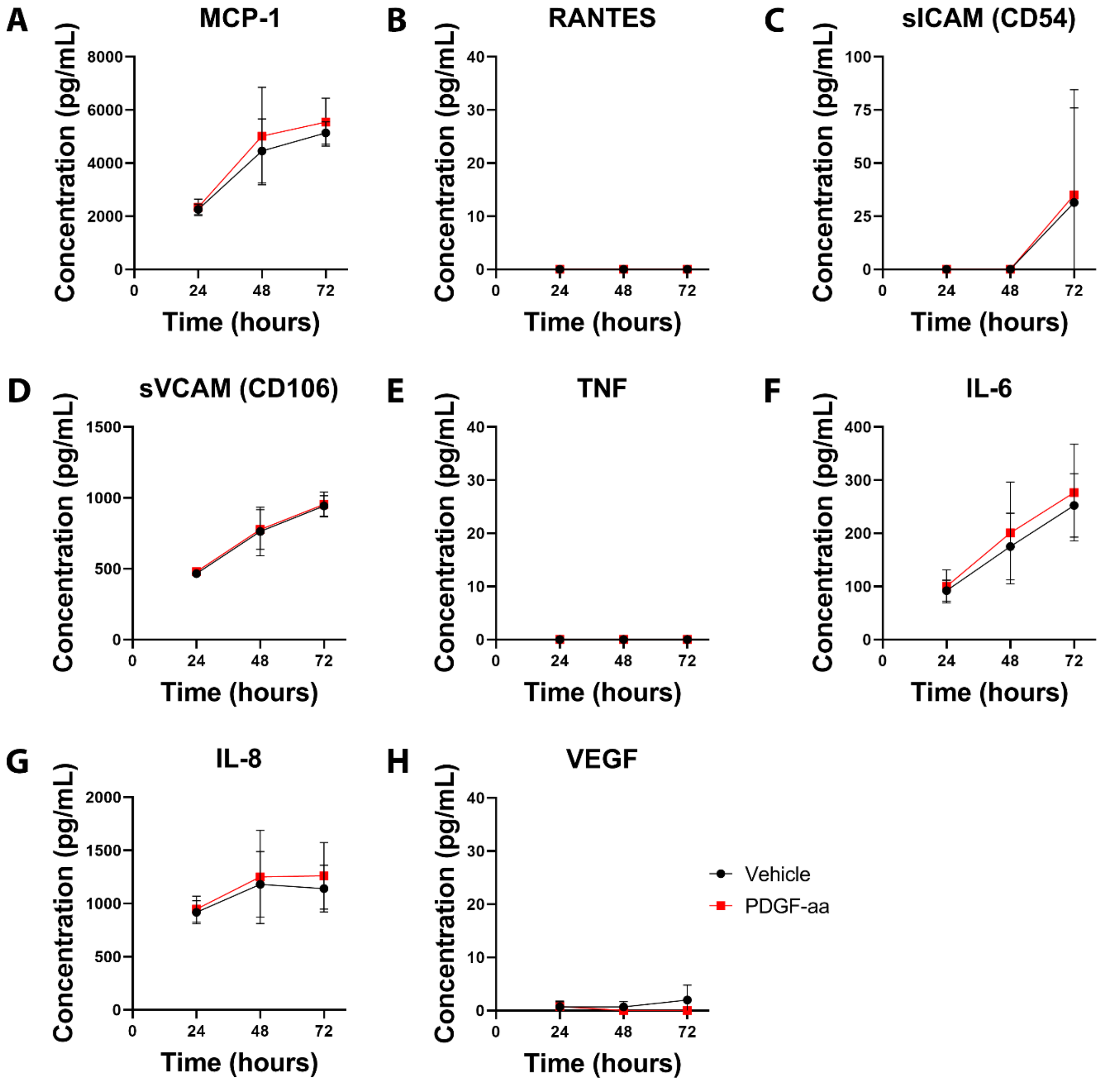


**Supplemental figure 7: Primary pericytes do not change secretion of inflammatory factors in response to PDGF-aa.** Pericyte medium was sampled with replacement 24, 48, and 72 hours after treatment with either PDGF-aa or vehicle. The concentration of MCP-1 (A), RANTES (B), sICAM (C), sVCAM (D), TNF (E), IL-6 (F), IL-8 (G), and VEGF (H) was investigated in each media sample by cytometric bead array. The presented data is collected from two experimental repeats.

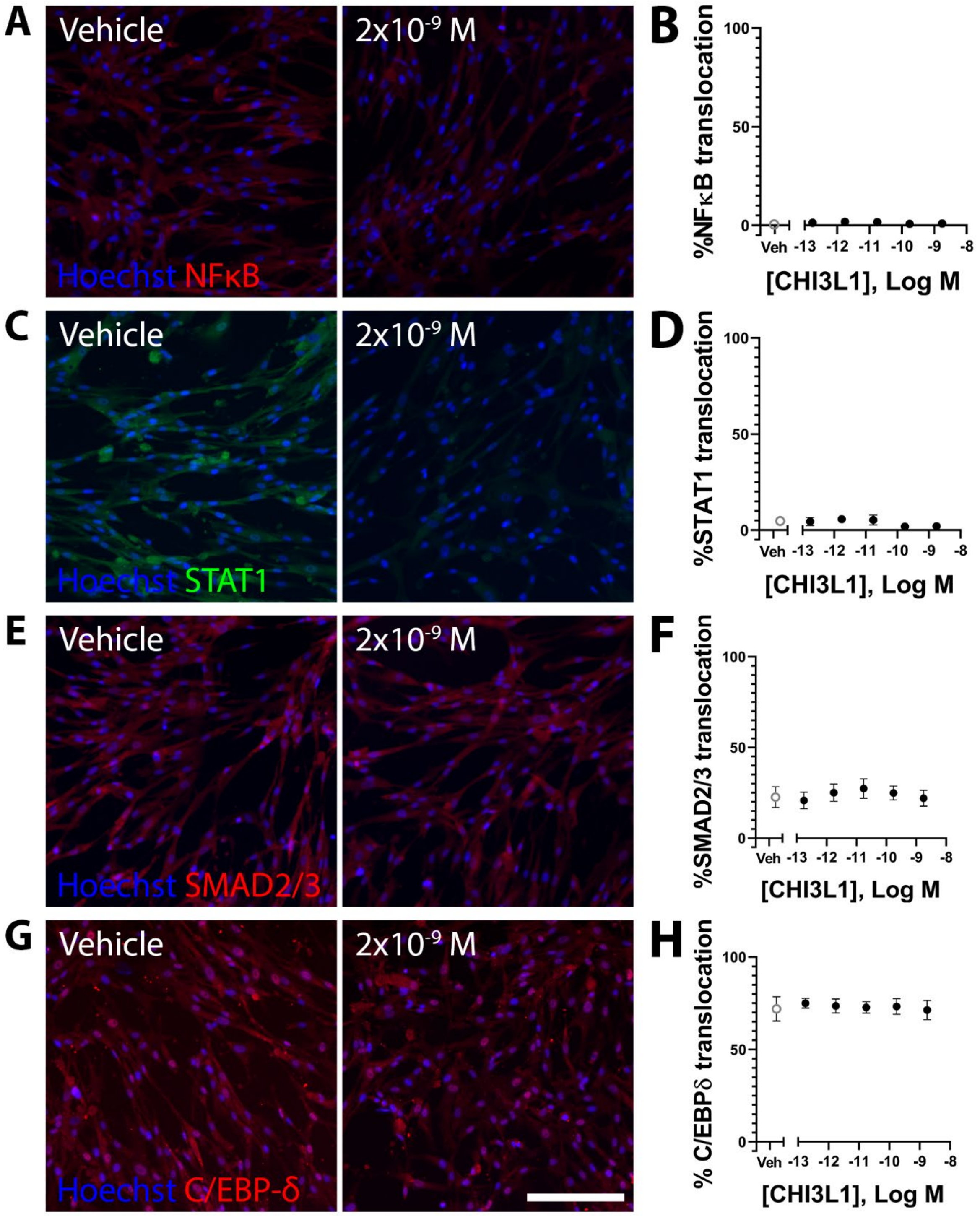


**Supplemental figure 8: CHI3L1 treatment does not cause translocation of NFκB, STAT1, SMAD2/3 or C/EBP-δ in primary pericytes.** Immunofluorescence images demonstrating no change in NFκB (A,B), STAT1 (C,D), SMAD2/3 (E,F), or C/EBP-δ (G,H) localisation in response to increasing concentrations of CHI3L1 in primary pericytes (only highest concentration shown). Images are quantified using MetaXpress to generate concentration-response curves (B,D,F,H) which demonstrate no CHI3L1-dependent change in signalling protein localisation. Data presented is one representative experiment of two experimental repeats. Scale bar = 200μm for all images.

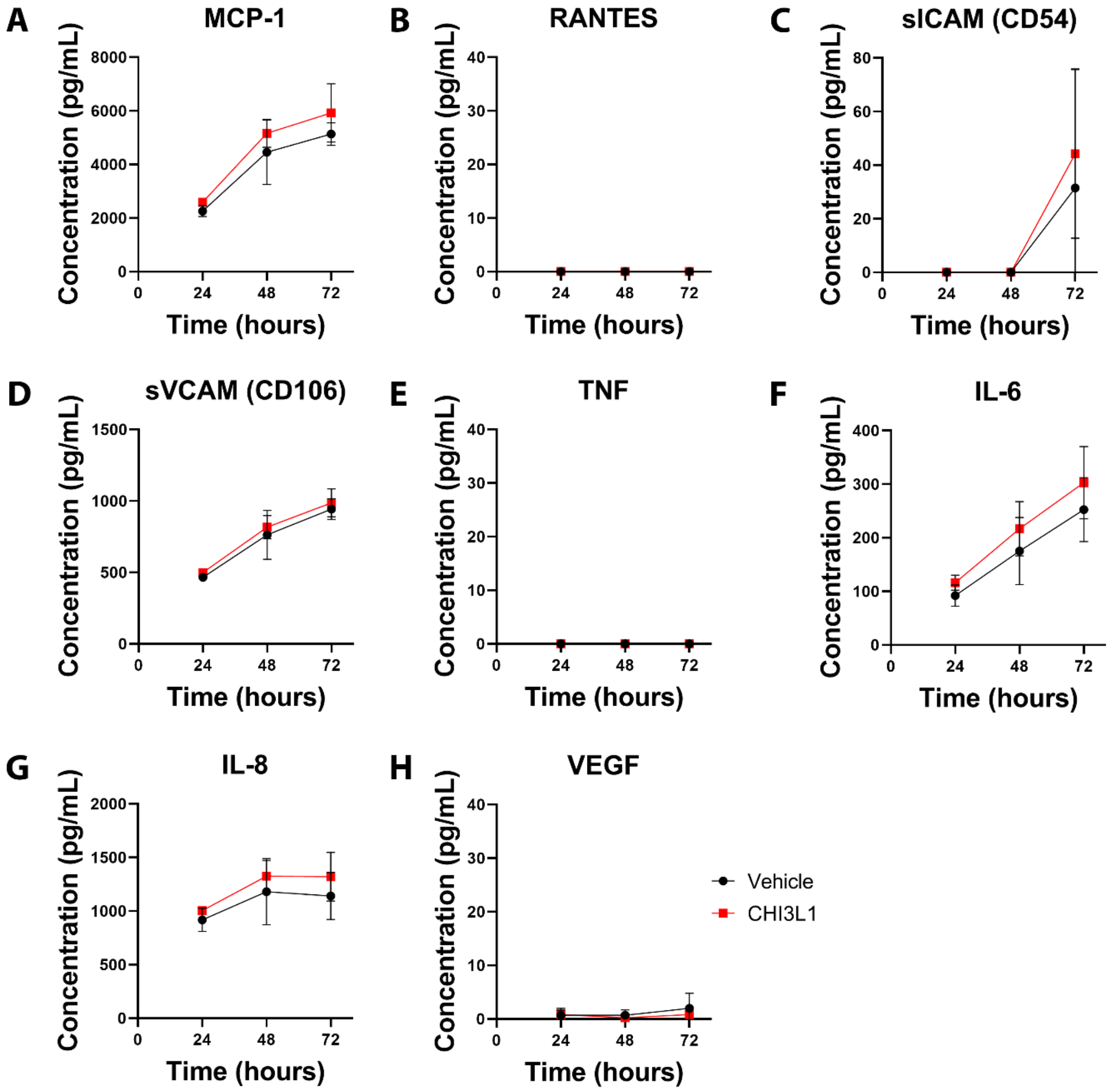


**Supplemental figure 9: Primary pericytes do not change secretion of inflammatory factors in response to CHI3L1.** Pericyte medium was sampled with replacement 24, 48, and 72 hours after treatment with either CHI3L1 or vehicle. The concentration of MCP-1 (A), RANTES (B), sICAM (C), sVCAM (D), TNF (E), IL-6 (F), IL-8 (G), and VEGF (H) was investigated in each media sample by cytometric bead array. The presented data is collected from two experimental repeats.

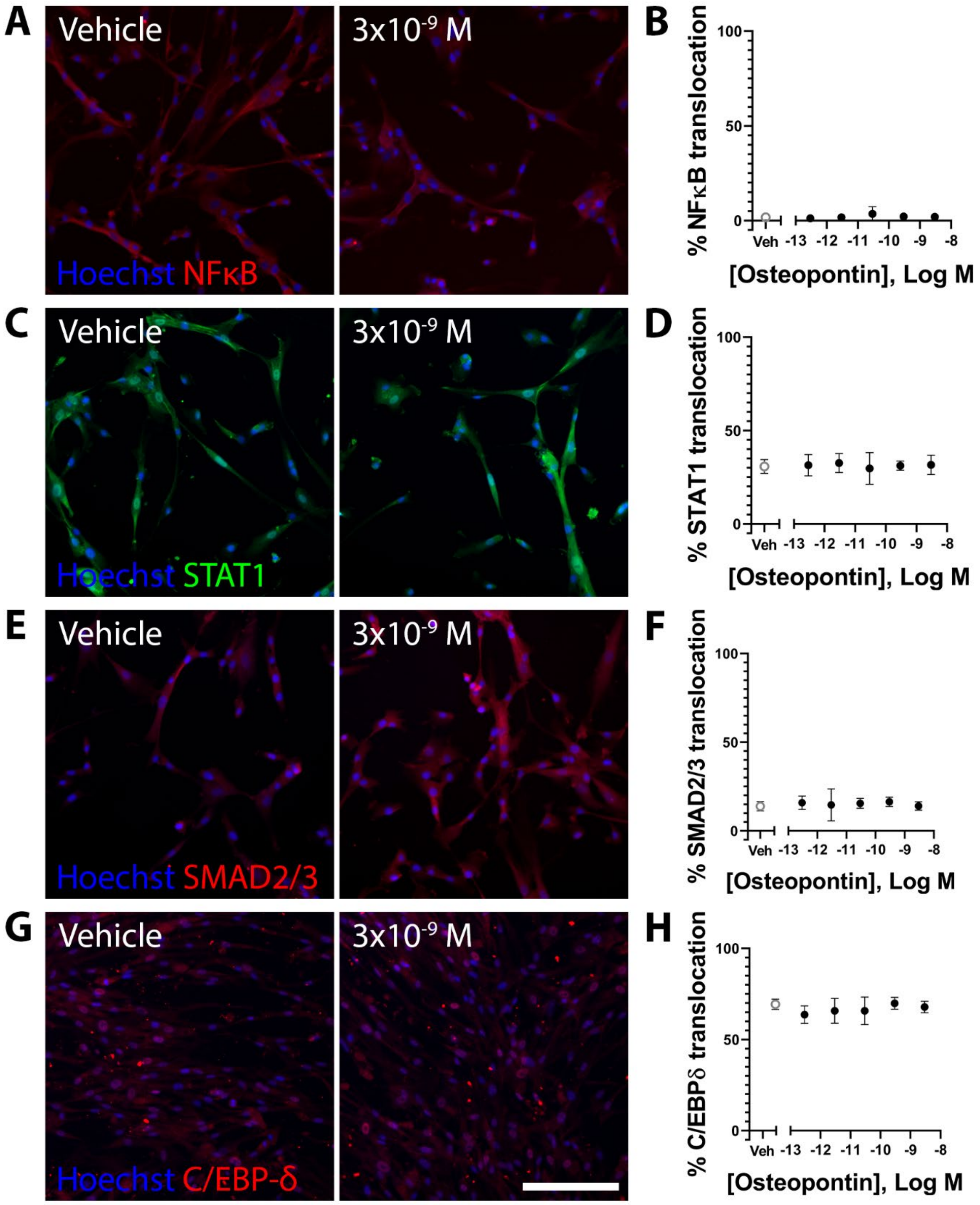


**Supplemental figure 10: Osteopontin treatment does not cause translocation of NFκB, STAT1, SMAD2/3 or C/EBP-δ in primary pericytes.** Immunofluorescence images demonstrating no change in NFκB (A,B), STAT1 (C,D), SMAD2/3 (E,F), or C/EBP-δ (G,H) localisation in response to increasing concentrations of osteopontin in primary pericytes (only highest concentration shown). Images are quantified using MetaXpress to generate concentration-response curves (B,D,F,H) which demonstrate no osteopontin-dependent change in signalling protein localisation. Data presented is one representative experiment of three experimental repeats. Scale bar = 200µm for all images.

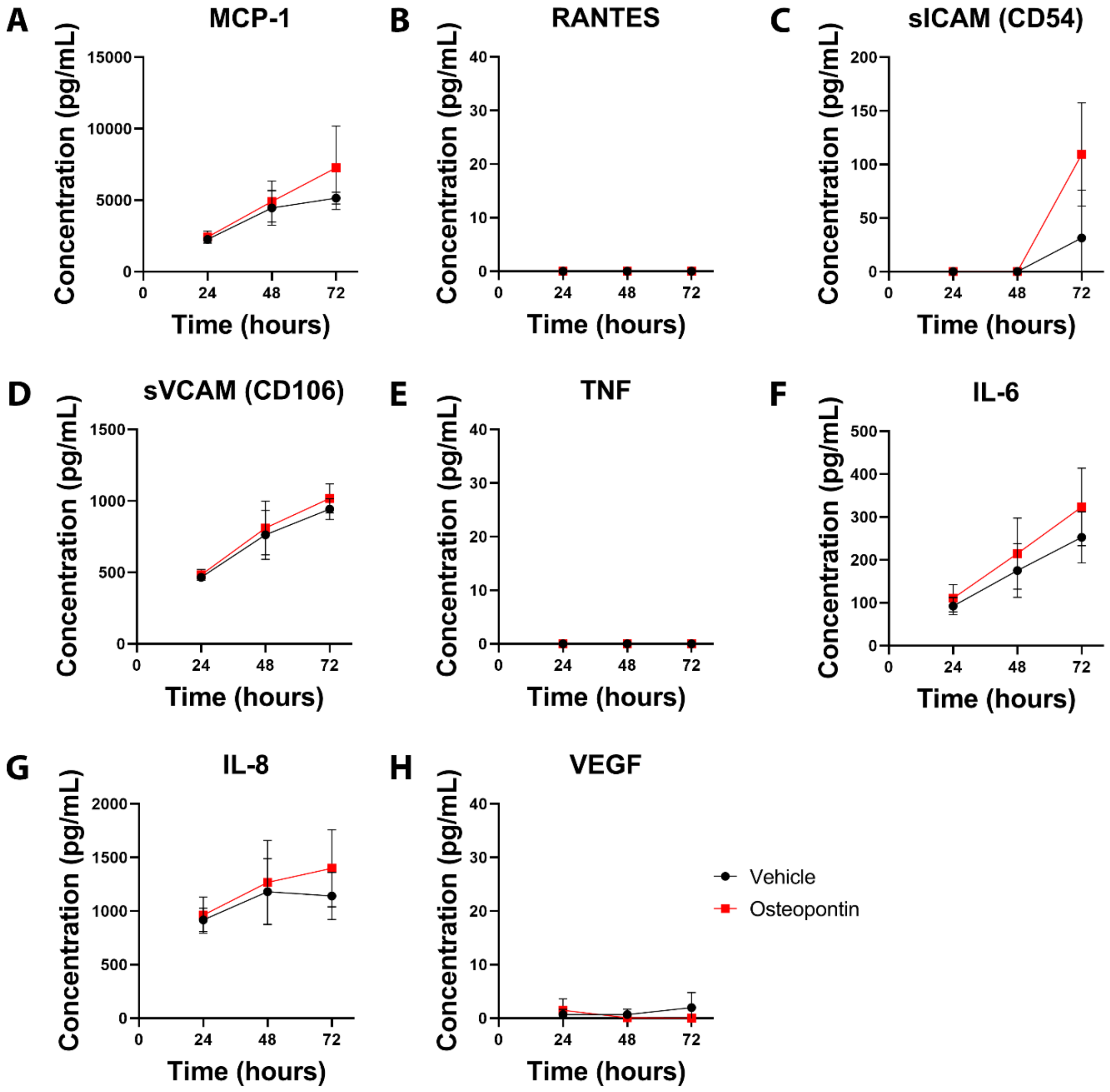


**Supplemental figure 11: Primary pericytes do not change secretion of inflammatory factors in response to osteopontin.** Pericyte medium was sampled with replacement 24, 48, and 72 hours after treatment with either osteopontin or vehicle. The concentration of MCP-1 (A), RANTES (B), sICAM (C), sVCAM (D), TNF (E), IL-6 (F), IL-8 (G), and VEGF (H) was investigated in each media sample by cytometric bead array. The presented data is collected from two experimental repeats.

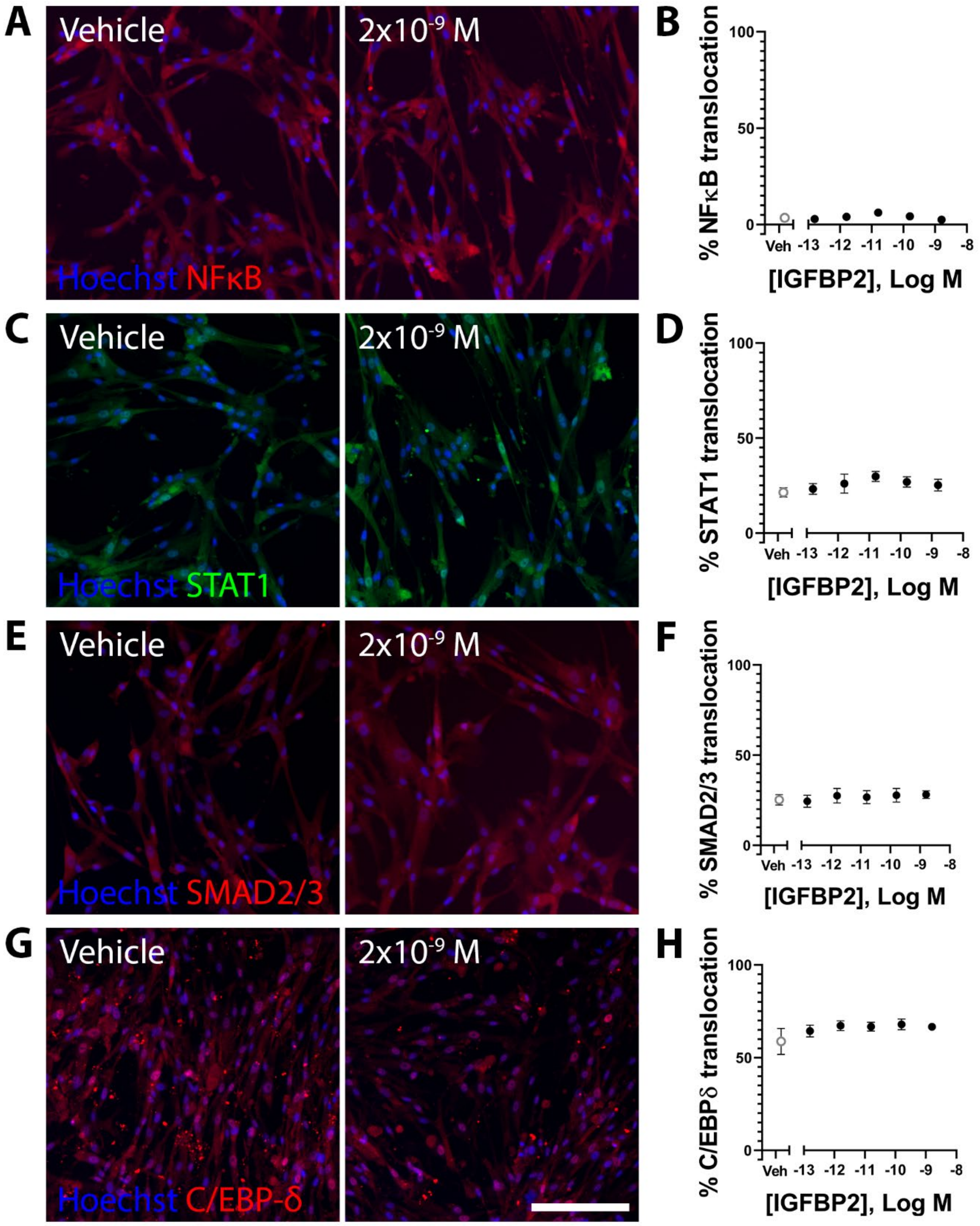


**Supplemental figure 12: IGFBP2 treatment does not cause translocation of NFκB, STAT1, SMAD2/3 or C/EBP-δ in primary pericytes.** Immunofluorescence images demonstrating no change in NFκB (A,B), STAT1 (C,D), SMAD2/3 (E,F), or C/EBP-δ (G,H) localisation in response to increasing concentrations of IGFBP2 in primary pericytes (only highest concentration shown). Images are quantified using MetaXpress to generate concentration-response curves (B,D,F,H) which demonstrate no IGFBP2-dependent change in signalling protein localisation. Data presented is one representative experiment of two experimental repeats. Scale bar = 200µm for all images.

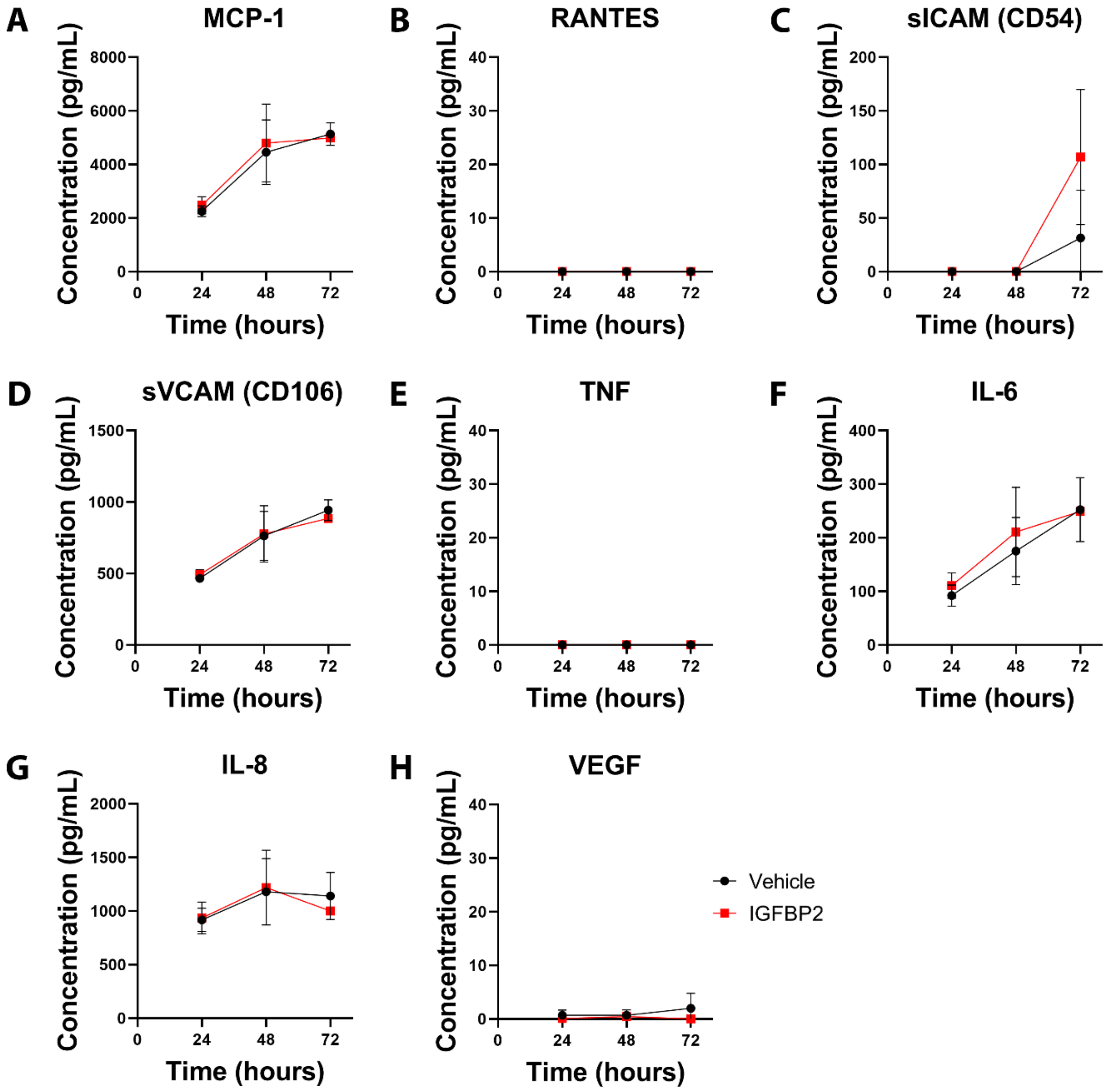


**Supplemental figure 13: Primary pericytes do not change secretion of inflammatory factors in response to IGFBP2.** Pericyte medium was sampled with replacement 24, 48, and 72 hours after treatment with either IGFBP2 or vehicle. The concentration of MCP-1 (A), RANTES (B), sICAM (C), sVCAM (D), TNF (E), IL-6 (F), IL-8 (G), and VEGF (H) was investigated in each media sample by cytometric bead array. The presented data is collected from two experimental repeats.

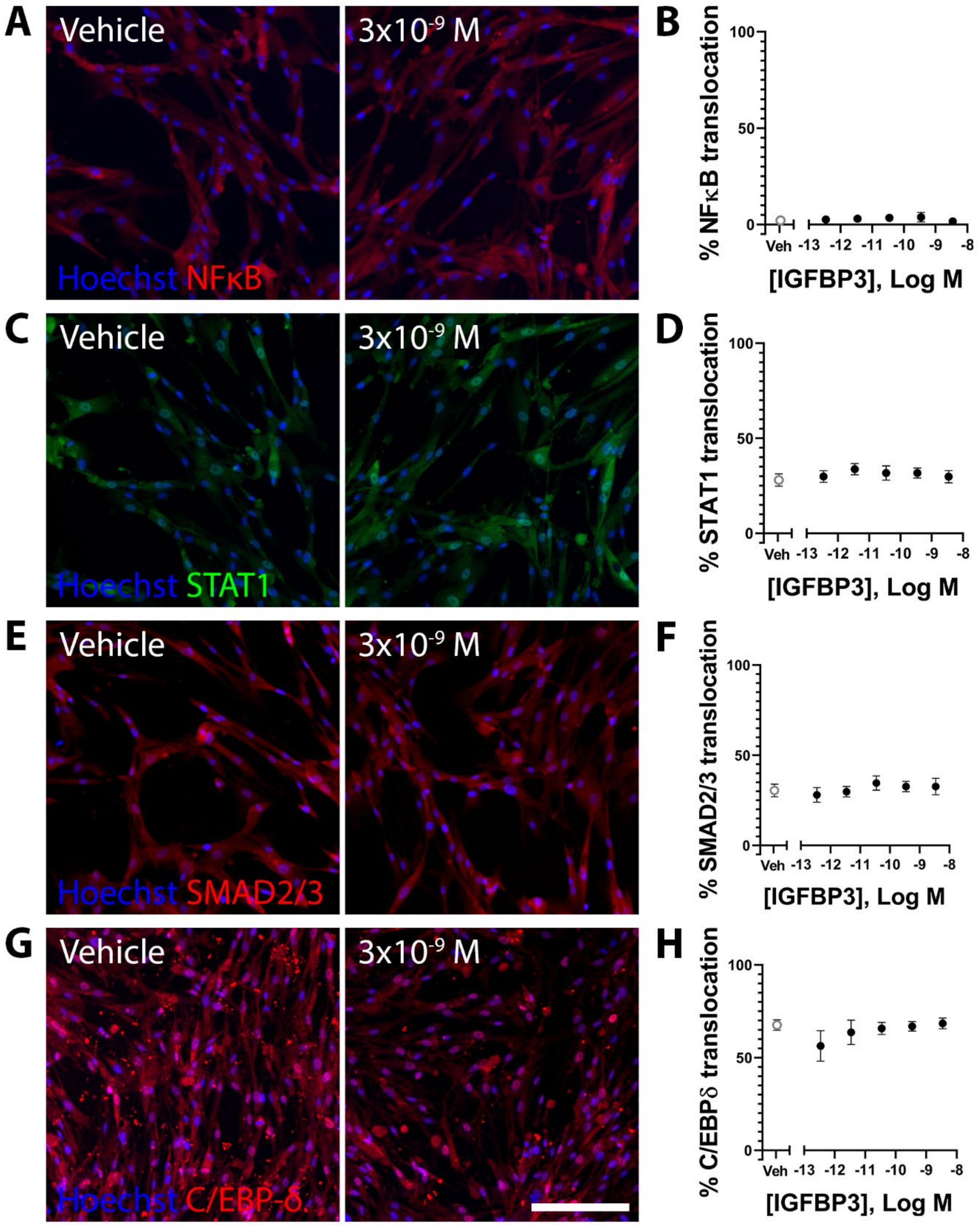


**Supplemental figure 14: IGFBP3 treatment does not cause translocation of NFκB, STAT1, SMAD2/3 or C/EBP-δ in primary pericytes.** Immunofluorescence images demonstrating no change in NFκB (A,B), STAT1 (C,D), SMAD2/3 (E,F), or C/EBP-δ (G,H) localisation in response to increasing concentrations of IGFBP3 in primary pericytes (only highest concentration shown). Images are quantified using MetaXpress to generate concentration-response curves (B,D,F,H) which demonstrate no IGFBP3-dependent change in signalling protein localisation. Data presented is one representative experiment of two experimental repeats. Scale bar = 200µm for all images.

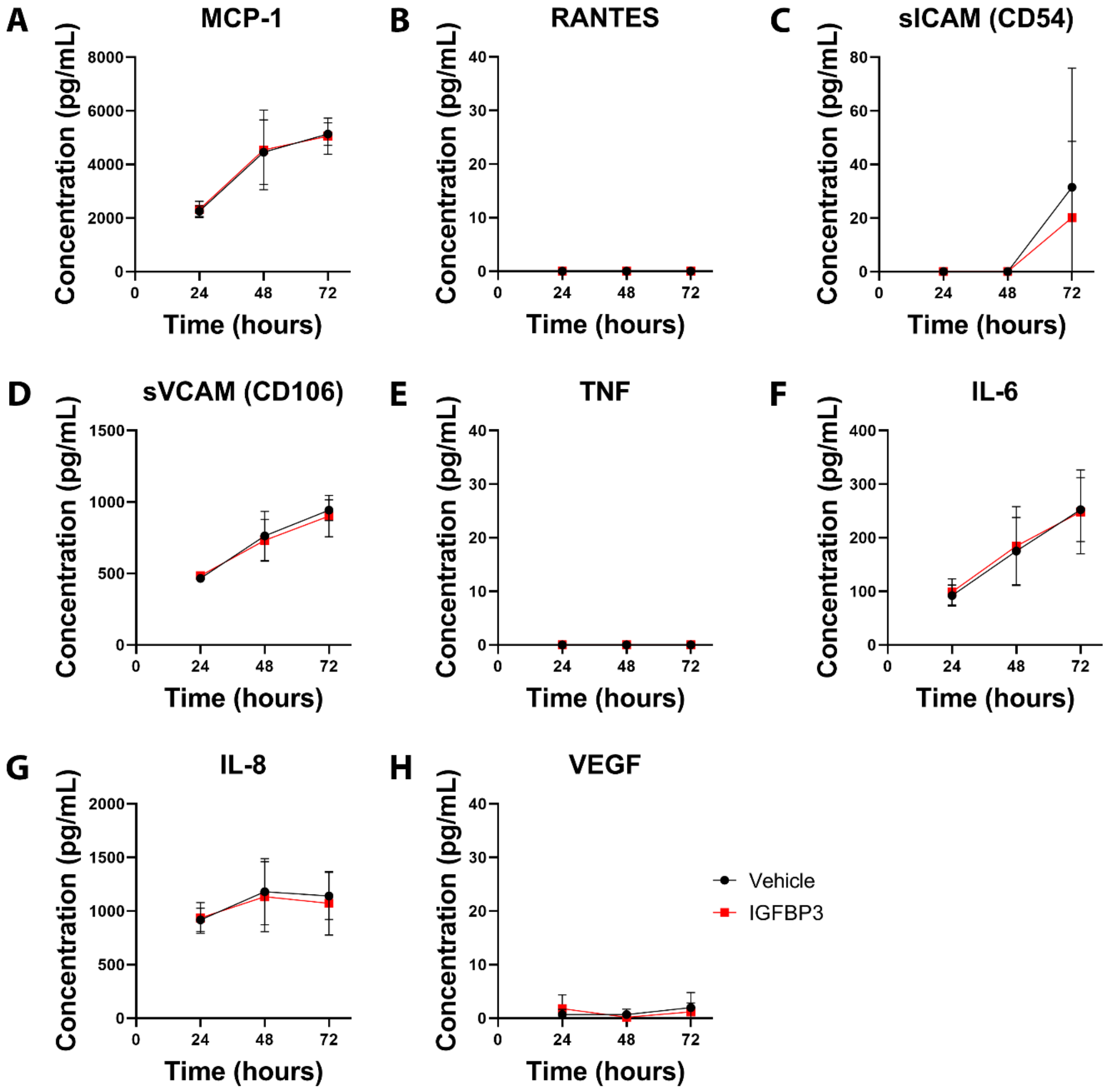


**Supplemental figure 15: Primary pericytes do not change secretion of inflammatory factors in response to IGFBP3.** Pericyte medium was sampled with replacement 24, 48, and 72 hours after treatment with either IGFBP3 or vehicle. The concentration of MCP-1 (A), RANTES (B), sICAM (C), sVCAM (D), TNF (E), IL-6 (F), IL-8 (G), and VEGF (H) was investigated in each media sample by cytometric bead array. The presented data is collected from two experimental repeats.

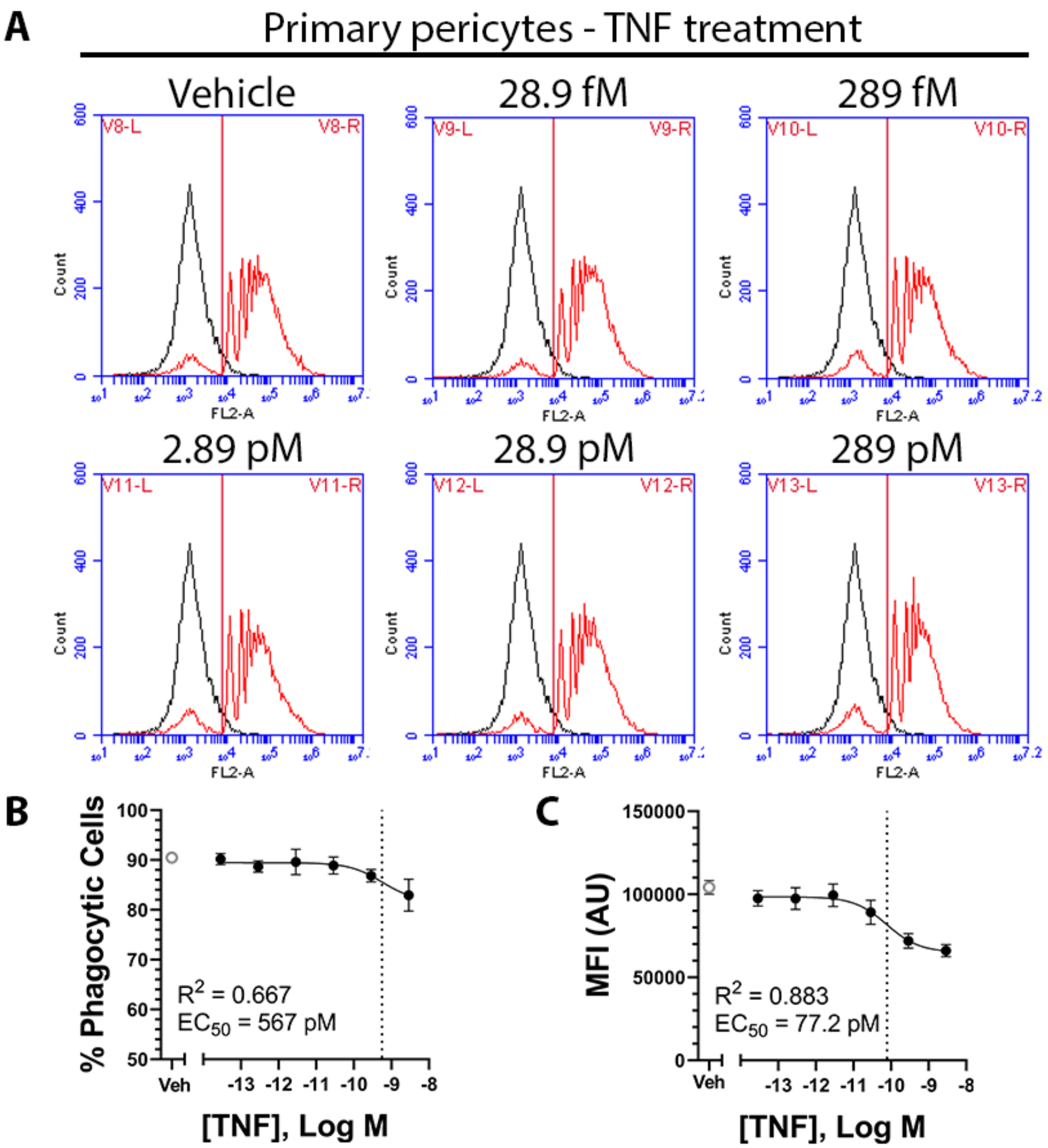


**Supplemental figure 16: TNF reduces the percentage of phagocytic cells and amount of phagocytosis in primary pericytes in a concentration-dependant manner.** (A) Fluorescence histo-plots demonstrating the distribution of fluorescent phagocytosis beads after treatment for 72 hours with increasing concentrations of TNF (red, with cell auto-fluorescence present in black). Beads are present for the last 24 hours of this treatment period. Concentration-response graphs are generated showing the percentage of phagocytic cells (B) and the mean fluorescent intensity (MFI) of phagocytic cells (C).

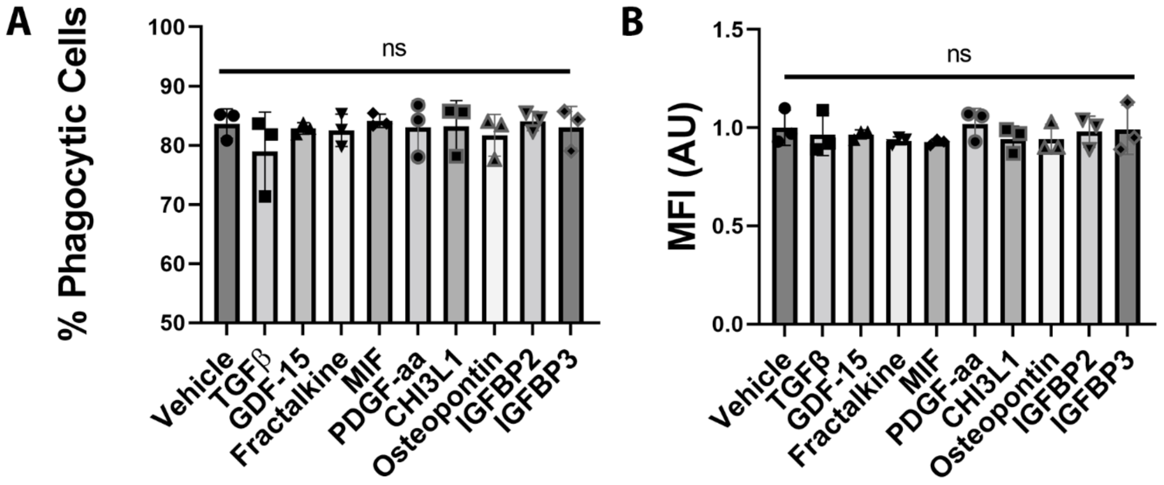


**Supplemental figure 17: TGFβ, GDF-15, fractalkine, MIF, PDGF-aa, CHI3L1, osteopontin IGFBP2 and IGFBP3 do not reduce phagocytosis in primary pericytes.**
Primary pericytes are treated with either vehicle, TGFβ (200pM), GDF-15 (8.13nM), fractalkine (588pM), MIF (333pM), PDGF-aa (175pM), osteopontin (2.97nM), IGFBP2 (1.59nM), or IGFBP3 (3.47nM) for 72 hours, with fluorescent beads present for the final 24 hours. Quantification of flow cytometry data was generated using the gating strategy described in Chapter 4, Figure 18, and shows no change in the percentage of phagocytic cells (A) and the mean fluorescent intensity (MFI) of phagocytic cells (B) between vehicle and any treatment. The presented data is collected from three experimental repeats. The significance of each treatment compared to vehicle was determined using a one-way ANOVA with multiple comparisons.

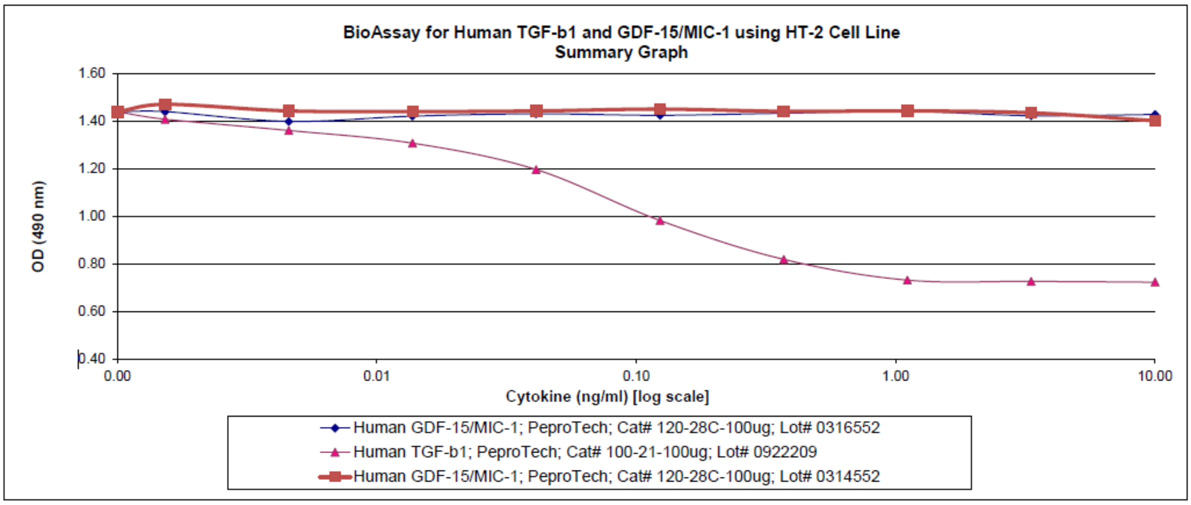


**Supplemental figure 18: Peprotech GDF-15 used in this study does not contain TGFβ.** A functional bioassay demonstrating no TGFβ response from the GDF-15 lot used in this study. Data was provided by Peprotech.

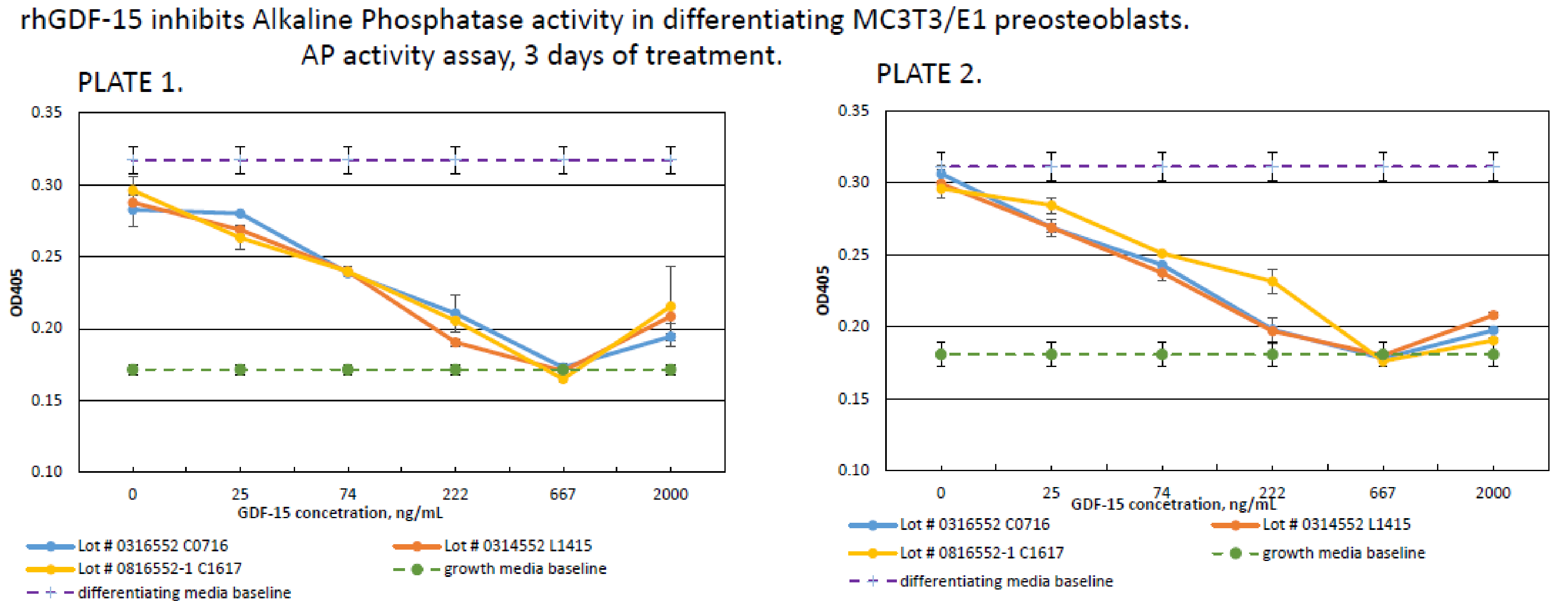


**Supplemental figure 19: Peprotech GDF-15 possess functional activity.** A functional bioassay demonstrating a GDF-15 response from the GDF-15 lot used in this study. Data was provided by Peprotech.
